# Switches in Orientation Coding by Mouse Primary Visual Cortex Neurons Depend on Stimulus Predictability

**DOI:** 10.1101/2025.06.29.662227

**Authors:** Jeroen M. van Daatselaar, Tom Sikkens, Mariel A. Muller, Cyriel M. A. Pennartz, Umberto Olcese, Conrado A. Bosman

**Affiliations:** Swammerdam Institute for Life Sciences, University of Amsterdam, Science Park 904, Amsterdam, 1098 XH, The Netherlands

## Abstract

How the brain processes a stimulus depends on contextual factors, such as whether it is predictable or surprising. While this process has been partially characterized using EEG and fMRI, and invasive cellular approaches, the microcircuit mechanisms responsible for comparing sensory input and expectations remain poorly understood. Here, we combined layer-resolved recordings of single-unit activity and local field potentials (LFP) in mouse primary visual cortex (V1) with a visual oddball paradigm using oriented gratings. Both event-related potentials and firing rates exhibited distinct temporal components across cortical layers and trial types. We identified robust stimulus-specific adaptation (SSA), mismatch negativity (MMN), and deviant detection (DD) contrast at both early and late epochs. Surprisingly, a substantial subset of excitatory neurons exhibited complete reversals of orientation preference (“preference switches”) between standard and deviant trials, rather than only changes in selectivity. These preference-switching cells were primarily observed in layers 2/3 and 5/6 and contributed as much information to stimulus decoding as stably tuned neurons. Our findings demonstrate that context-dependent flexibility in feature preference is an integral part of predictive coding in V1 and challenge the notion of fixed stimulus representations at the single-neuron level.

## Introduction

Unexpected events influence us in ways that predictable ones do not, as illustrated when a routine is disrupted or when the timing, occurrence, or characteristics of an event violate expectations. These deviations highlight the brain’s reliance on predictability, as it continuously extracts regularities from the environment to form predictions and compare them against incoming sensory input^1–6^. This ability to anticipate events and detect deviations is crucial for adaptive behavior, enabling rapid responses to unexpected changes and facilitating learning from them.

The predictive coding (PC) framework is well suited to explore neural processes involved in deviance detection^1–3,6,7^. Classic PC posits that each anatomical level in the cortical hierarchy compares incoming sensory input to predictions generated by an internal model of the world^2–4,6,8–10^. With experience, regularities in past events are translated into predictions of upcoming sensory input, which in turn serve to ‘explain away’ the regularity. This process can partially account for stimulus-specific adaptation (SSA)^10–14^, where responses to prediction-affirming stimuli gradually decrease. In contrast, unpredicted aspects of sensory input are propagated up the cortical hierarchy as ‘prediction error,’ potentially explaining the exaggerated neural response to surprising events – an effect known as deviance detection (DD)^10,11,15–23^. SSA and DD are thought to be components of the pattern of mismatch negativity (MMN) responses, which has been extensively studied using the oddball paradigm^9,10,24^. More recently, the inclusion of control conditions has allowed MMN to be experimentally partitioned into SSA and DD, both at the level of single cells^15,17,22,23,25–28^ and aggregate neuronal responses such as Local Field Potential (LFPs) and multi-unit activity (MUA)^15,26,27,29^. PC thus offers an interpretation of MMN, SSA, and DD, which coheres well with empirical observations^6,7,9,10,30^. However, while PC provides a clear algorithmic explanation, precisely how it is implemented at the cortical microcircuit level remains unclear.

Recent studies, informed by PC theory, have begun to unravel the neural mechanisms underlying DD^1,27,31,32^. These studies suggest that DD arises from the interplay of feedforward and feedback inputs to cortical circuits, with prediction-specific patterns of feedback activity interacting with sensory-driven feedforward inputs in supragranular layer 2/3 (L2/3) of primary sensory cortices, in line with computational models^4,33,34^. However, current models are primarily limited to L2/3 and offer less evidence on the role of granular (L4) and infragranular layers (L5/6), despite evidence of their distinct contributions to cortical processing^4,35^.

Many previous studies have treated the neural response as a single event, overlooking the temporal complexity of deviant detection. Yet DD comprises at least two distinct phases: an early (50-200 ms) component linked to feedforward processing, and a later (200-400 ms) stage often associated with feedback and integration^22,25,36–38^. How these phases map onto specific cortical layers and circuit mechanisms remains unclear. Disentangling the interplay between feedforward and feedback inputs across layers and time is therefore essential for understanding deviance detection.

Given the influence of top-down signals on cortical dynamics, a key question is whether they modulate not just the gain or timing of responses, but also the content of sensory representations. Orientation selectivity, a fundamental property of V1 neurons, is classically viewed as stable under normal conditions^39–42^. However, emerging evidence shows that top-down signals, especially those related to prediction and expectation, can modulate orientation selectivity in V1^43–45^. Whether these influences can induce context-dependent shifts in tuning, rather than merely adjusting response strength, remains unclear. Since orientation selectivity is generally thought to be stable, understanding if and how predictability or surprise can dynamically alter a neuron’s preferred orientation is crucial for clarifying how unexpected sensory events reshape cortical representations^1,4,6,33^.

Here, we addressed these questions by combining laminar recordings of local field potentials (LFPs) and single-unit activity in mouse V1 with a visual oddball paradigm using horizontal and vertical gratings. We systematically quantified SSA, MMN, and DD effects at both the population and single-neuron level across cortical layers and temporal epochs. Surprisingly, we observed not only charges in response strength but also complete reversals of orientation preference (“preference switches”) in a substantial subset of excitatory neurons, primarily in extragranular layers. We further show that these context-dependent preference switches do not impair the ability of V1 population to encode stimulus orientation, but rather coexist with robust stimulus-specific information. These findings challenge the view of fixed tuning in early sensory cortex and suggest that predictive context can transiently reshape feature representations at the single-neuron level.

## Results

### Neural Responses in the Oddball Task

To explore how V1 processes unexpected and expected visual stimuli, we recorded single-neuron activity and local field potentials in awake, head-fixed mice while they were shown drifting gratings at varying orientations (see *Methods*). Mice were acutely implanted with a two-shank multi-channel electrode array spanning all cortical layers in V1 and passively presented with a visual oddball task, along with a many-standards control task (Figure 1A). In the oddball task, deviant stimuli consisted of a 90° orientation change relative to the repeated standard stimuli. Trial types from both tasks were labeled as “many-standards control” (ctr), “standard” (std), or “deviant” (dev). The standard stimulus, being the last in a sequence of identical stimuli, provides an event that conforms to a predictable pattern on a local time scale, within the immediate temporal structure of the sequence. Every 6-10 trials, this sequence was interrupted by a deviant stimulus, breaking the repetition with an unexpected event. The control stimuli were physically identical to the deviant and occurred with equal probability, but were embedded within an unpredictable sequence of stimuli, ensuring that no single stimulus could be anticipated based on prior trials.

**Figure 1.**
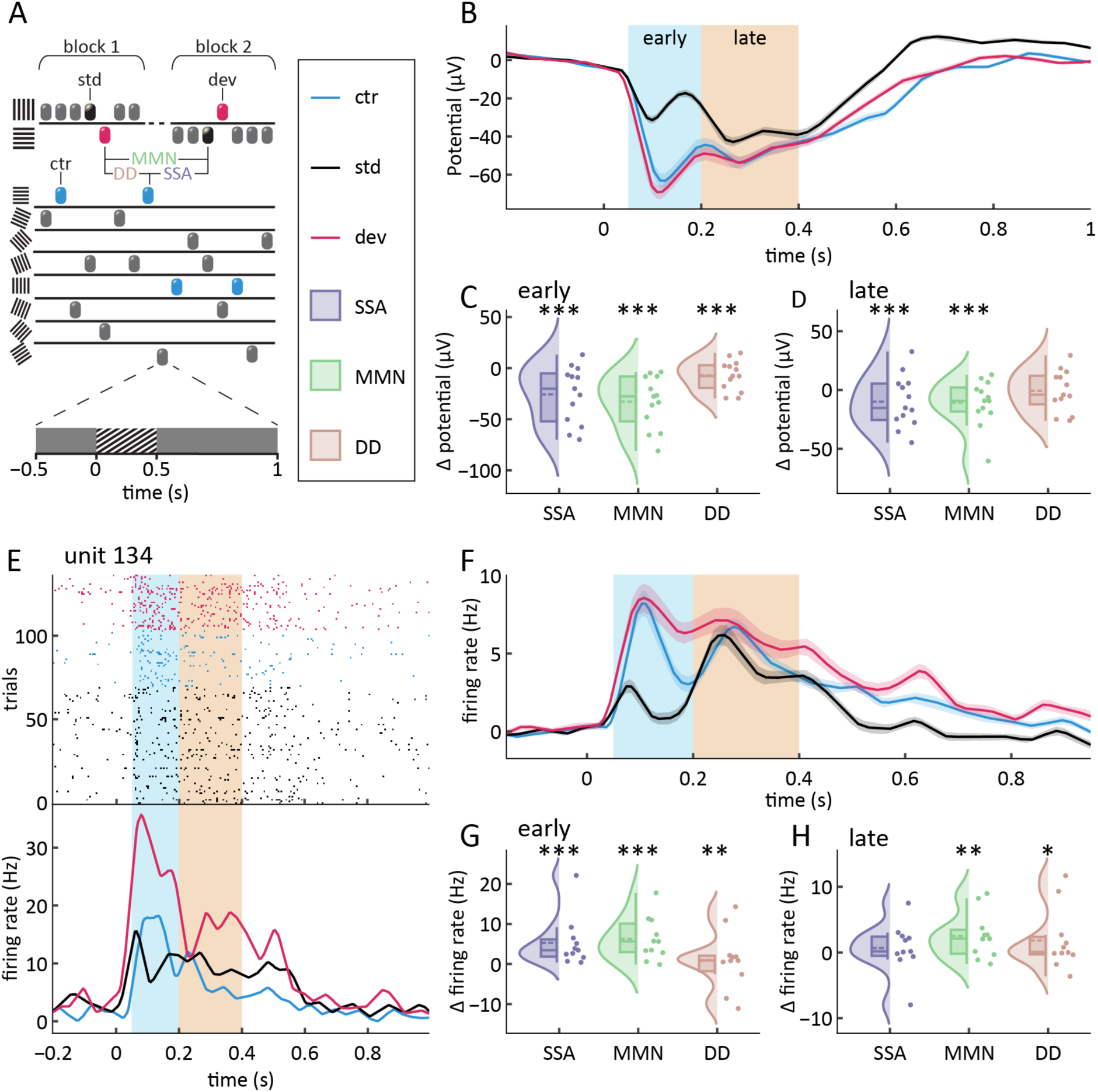
Stimulus context modulates early and late local field potential and neuronal responses. (A) Schematic of task design. The upper portion of the diagram depicts a classic counterbalanced oddball design. The lower part of the diagram shows the many-standards control task. Black, blue, and magenta icons indicate standard, deviant, and control trials, respectively. Trial contrasts SSA, MMN, and DD (lavender, green, and rose-tan, respectively) are computed by subtracting trial types with identical stimuli and subsequent averaging across stimulus types. Legend: std, standard; dev, deviant; ctr, control; SSA, stimulus-specific adaptation; MMN, mismatch negativity; DD, deviance detection. (B) Average ERP of all recorded channels after baseline correction. Marked areas indicate time windows designated as ‘early’ (50-200 ms) and ‘late’ (200-400 ms) (n = 252). (C and D) Average ERP trial type effects for (C) early and (D) late latencies. Each dot represents the average of one recording session. Difference potentials (Δ-potentials) are derived from the SSA, MMN, and DD trial contrasts. (E) Raster plot and corresponding peri-stimulus time histogram (PSTH) of one typical example neuron. Colors of traces are as in (A). (F) Average PSTH of all recorded single neurons after baseline correction (n = 121). (G and H) Average firing rate trial type effects per recording session for (C) early and (D) late latencies. Changes in firing rates (Δ-firing rates) are computed based on the SSA, MMN, and DD trial contrasts. Line plots show mean ± SEM, smoothed at 100 ms using a Tukey window. Raincloud plots show all individual data points, smoothed distribution, and boxplot with whiskers (outlier limits), interquartile range, box (quartile 1 and 3 upper limits), median (solid line), and mean (dashed line). *p < 0.05, **p < 0.01, ***p < 0.001 indicate significance from zero (i.e., no change in firing rate), based on a linear mixed-effects model analysis (see Methods and Supplementary Tables 1-4).

To quantify responses to these conditions, we computed three contrasts: stimulus-specific adaptation (SSA, std – ctr), mismatch negativity (MMN, dev – std), and deviant detection (DD, dev – ctr) (Figure 1A). Most statistical comparisons were assessed using linear mixed-effect models, which accounted for within-subject variability by including recording session, electrode channel, neuron identity, or layer specificity as random effects. These analyses are reported in Supplementary Tables 1-17. Alternative analyses involving permutation tests, decoding-based comparisons, and proportion tests were used where appropriate and are described in the corresponding Results sections and Supplementary Tables 18-25 (see *Methods* for details).

To replicate prior findings on oddball responses, we pooled data from all intracortical electrodes (N = 252) within each session (N = 13) and computed event-related potentials (ERPs) from the averaged LFP traces across channels. Clear ERP responses consistently emerged with two main temporal components (Figure 1B), matching previously described processing stages^22,25,36,37^. We then calculated response amplitudes by averaging ERPs in temporal windows (hereafter referred to as early and late epochs, 50-200 ms and 200-400 ms, respectively) aligned with the two observed response components. During the early period, all three contrasts, SSA, MMN, and DD, showed significant differences from zero (SSA: *p* < 0.001; MMN: *p* < 0.001; DD: *p* = 0.0002; Figure 1C; Supplementary Table 1). During the late period, ERP traces for different trial types gradually converged to a similar amplitude, with SSA and MMN still showing significant differences from zero (SSA: *p* = 0.0199; MMN: *p* = 0.0204), while DD no longer reached significance (DD: *p* = 0.4752; Figure 1D; Supplementary Table 2). These findings align with prior observations in rodent and primate cortices, including sensory and associative areas, regarding response amplitudes for control, standard, and deviant stimuli, and confirm the existence of early and late response components^15,22,25,36,38^.

We next examined single-neuron responses. From 958 recorded units in V1, we first excluded multi-unit clusters and non-cortical units, yielding 151 cortical neurons. We then focused on putative excitatory neurons, identified via their action potential waveform characteristics (see *Methods*), resulting in a final sample of 141 neurons. Of these, 121 showed significant responsiveness to at least one combination of trial type and stimulus orientation and were included in the subsequent analyses. Responses were generally lowest for the standard stimulus, with firing rate peaks at similar latencies to those seen in ERPs (Figures 1E-F). SSA, MMN, and DD contrasts were significantly different from zero during early latency (*p* = 0.0047, *p* = 0.0083, and *p* = 0.0122, respectively; Figure 1G; Supplementary Table 3). In contrast, in the late window, only MMN and DD contrasts reached significance (*p* = 0.0118 and *p* = 0.0028, respectively), while SSA did not (*p* = 0.0675; Figure 1H; Supplementary Table 4). These results confirm a two-stage neural response pattern and indicate that V1 neurons in mice are sensitive to changes in stimulus predictability^23,28,38,46^.

### Layer-Specific Responses Reveal Early Supragranular and Late Infragranular Deviance Detection

Layer-specific LFP and spiking responses can be indicative of feedforward and feedback contributions to local neuronal activity^4,35,47^. To assess oddball responses separately for all cortical layers, we used current source density (CSD) and multi-unit activity (MUA) profiles to estimate the depth of each electrode relative to the cortical surface^48–50^ (Figure 2A-C; see Methods). Based on these profiles, channels from all sessions were grouped as supragranular (L2/3, N=96), granular (L4, N=26), and infragranular (L5/6, N=130).

**Figure 2.**
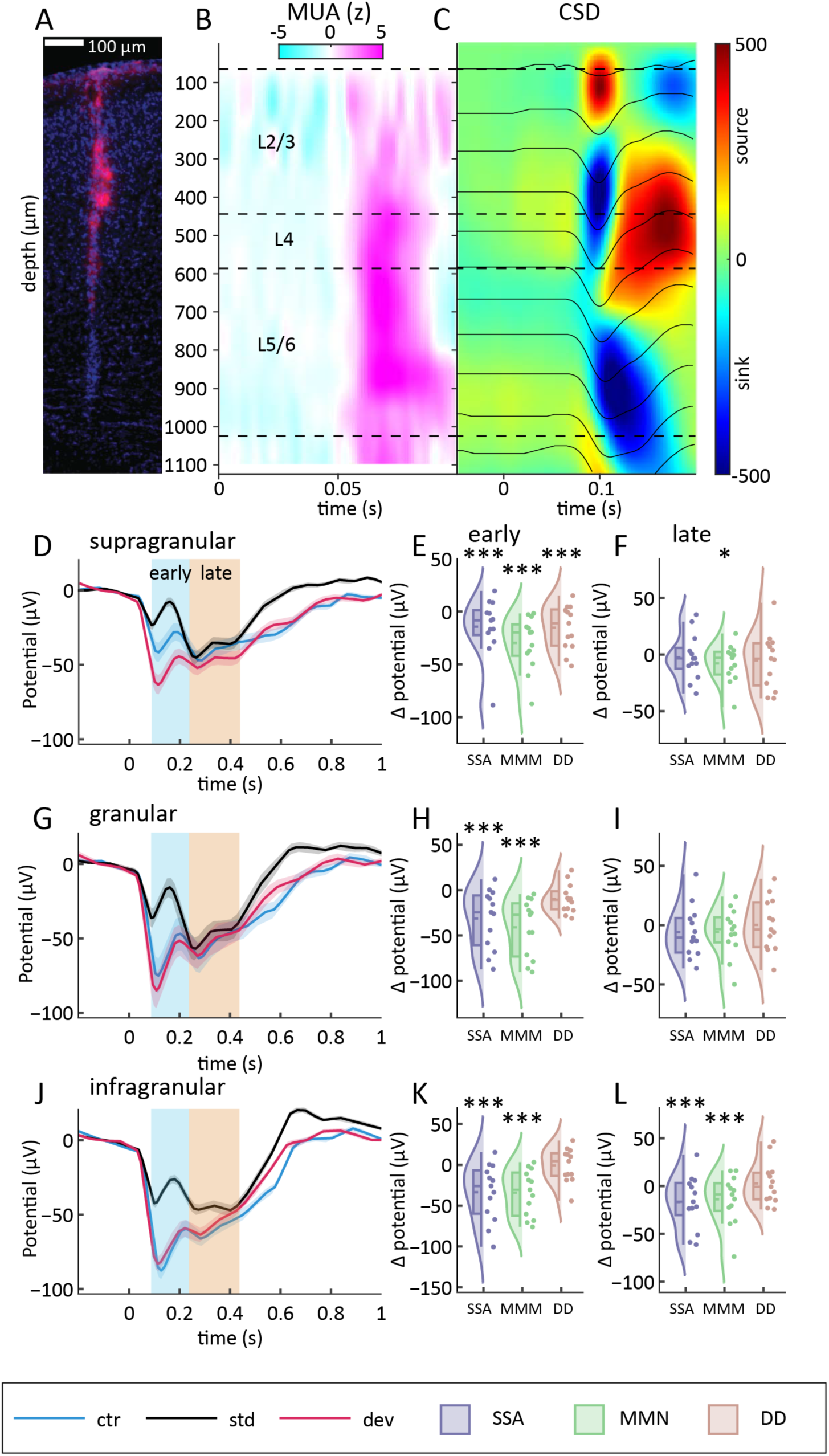
Early Supragranular and Late Infragranular ERP Effects During Visual Oddball Processing. (A-C) Example of indicators used to estimate probe penetration depth. (A) Staining of mouse V1. Magenta: DiI, purple: DAPI. Scale bar: 100 µm. (B) Multi-unit activity (MUA; arbitrary unit) along the cortical probe. Each channel of a probe (N = 16) was z-scored separately to facilitate interpretation. (C) Current-source density (CSD) plot with smoothed ERPs overlaid. Time zero (in seconds) marks stimulus onset. (D) Average ERP of all recordings for electrodes located in supragranular layers (L2/3) (n = 96). (E and F) Raincloud plots showing ERP trial type effects for (E) early and (F) late latencies. (G-H) As D-F, but for electrode sites located in granular layers (L4) (n = 26). (J-L) As D-F, but for electrode sites located in infragranular layers (L5/6) (n = 130). Plotting conventions as in Figure 1. *p < 0.05, **p < 0.01, ***p < 0.001 indicate significance from zero (i.e., no change in potential), based on linear mixed-effects model analysis (see Methods and Supplementary Tables 5-11).

We first analyzed ERP responses. The amplitude of control stimulus responses increased with cortical depth, both in absolute terms and relative to other trial types (see the vertical progression in Figures 2D, G, J; Supplementary Table 5). To facilitate direct comparisons across layers, y-axis limits were unified for all panels. Specifically, control response amplitudes were greater in granular compared to supragranular layers (*p* = 0.0022), in infragranular compared to granular layers (*p* = 0.0045), and in infragranular compared to supragranular layers (*p* = 0.0001).

Within each layer, standard stimuli consistently evoked the smallest early response amplitude (see trial-type differences in Figures 2D, G, J; Supplementary Tables 6-8). Both control and deviant stimuli produced significantly larger early responses compared to standard in supragranular (p = 0.0045 and p = 0.0087, respectively), granular (*p* = 0.0102 and *p* = 0.0136), and infragranular layers (*p* = 0.0031 and *p* = 0.0064). Taken together, these results indicate that trial type robustly influences early response amplitude across cortical layers.

To further characterize these effects, we quantified the key contrasts, SSA, MMN, and DD, within each layer. Both SSA and MMN responses were observed across all layers: for supragranular (*p* = 0.0045 for SSA, *p* = 0.0087 for MMN), granular (*p* = 0.0102 for SSA, *p* = 0.0136 for MMN), and infragranular layers (*p* = 0.0031 for SSA, *p* = 0.0064 for MMN). Early DD was significant only present in supragranular layers (*p* = 0.0117), but not in granular (*p* = 0.1902) or infragranular (*p* = 0.1091) layers (Figures 2E, K, H; Supplementary Tables 6-8).

During the late response phase, SSA emerged exclusively in infragranular layers *(p* = 0.0244), while granular (*p* = 0.0842) and supragranular (*p* = 0.1795) layers showed no significant SSA effect (Figures 2F, I, L; Supplementary Tables 9-11). In contrast, late response amplitudes in the granular layer showed no significant differences between trial types (*p* > 0.05 for all contrasts), suggesting that responses in this layer were primarily driven by bottom-up sensory input rather than contextual influences.

Next, we examined single-neuron activity across layers using the same grouping procedure. Neurons were classified as supragranular (N = 27), granular (N = 18), or infragranular (N = 76) (Figure S1). A power analysis indicated that statistical power was sufficient only for infragranular layers. During the early response epoch, MMN effects were observed across all layers (*p* = 0.0247, 0.0266, and 0.0109 for supragranular, granular, and infragranular neurons, respectively), whereas significant SSA was restricted to infragranular neurons (*p* = 0.034; *p* < 0.05 in other layer; Supplementary Tables 12-14). In the late response epoch, infragranular neurons exhibited both DD (*p* = 0.0165) and MMN (*p* = 0.0122) effects (Supplementary Tables 15-17), suggesting that deeper layers become selectively engaged in processing deviant stimuli at later stages.

These findings suggest a dynamic shift in the contributions of distinct cortical circuits to deviance processing across cortical layers and response stages. The early emergence of DD in supragranular layers may reflect an initial detection process that could involve top-down modulations, consistent with previous models^1,51,52^. However, the precise relationship between early supragranular DD effects and feedback input timing remains to be clarified, as feedback signals can reach the cortex rapidly in some contexts and are not exclusively associated with late processing windows. In contrast, the late onset of DD and MMN in infragranular layers indicates that deeper cortical circuits may integrate deviance-related signals over time^53^. Although our results do not pinpoint the involvement of feedforward or feedback pathways in specific layers or time windows, the distinct laminar and temporal patterns we observe highlight the complex circuitry underlying deviance processing in V1. To further characterize how V1 population dynamics contribute to this processing, we next examined activity patterns at single-cell population level.

### V1 Population Activity Shows Context-Dependent Changes in Orientation Encoding

For our population analyses, we split the data across nine conditions, corresponding to the different combinations of trial type (control, standard, deviant) and temporal epoch (baseline, early, late). We focused on two aspects of population activity: (1) the amount of stimulus-specific information represented in each condition, and (2) the extent to which population activity shifted as a function of trial type and epoch. To examine how much stimulus-specific information was present in each condition, we used a k-nearest neighbors (KNN) classifier to predict grating orientation from the spiking activity of all identified cortical excitatory neurons (N = 141; see *Methods*). Before stimulus onset, decoder accuracy was significantly above chance for standard (*p* = 0.014) and deviant (*p* = 0.004), but not for control (p = 0.56; Figure 3A; Supplementary Table 18). This is consistent with prior reports showing that regularity in the stimulus sequence biases population- and single-cell activity in a stimulus-specific manner prior to stimulus onset^27,28,46,54,55^. Upon stimulus presentation, decoding accuracy for the deviant condition increased during the early epoch (*p* < 0.001), while improvements for standard (*p* = 0.21) and control (*p* = 0.12) conditions only reached significance during the late epoch (standard: *p* = 0.009; control: *p* = 0.022; Figure 3B-C; Supplementary Table 18). These results demonstrate that stimulus sequence regularity induces stimulus-specific activity even before stimulus onset, and that violations of regularity (mismatch) enhance stimulus-specific representations shortly after stimulus presentation.

**Figure 3.**
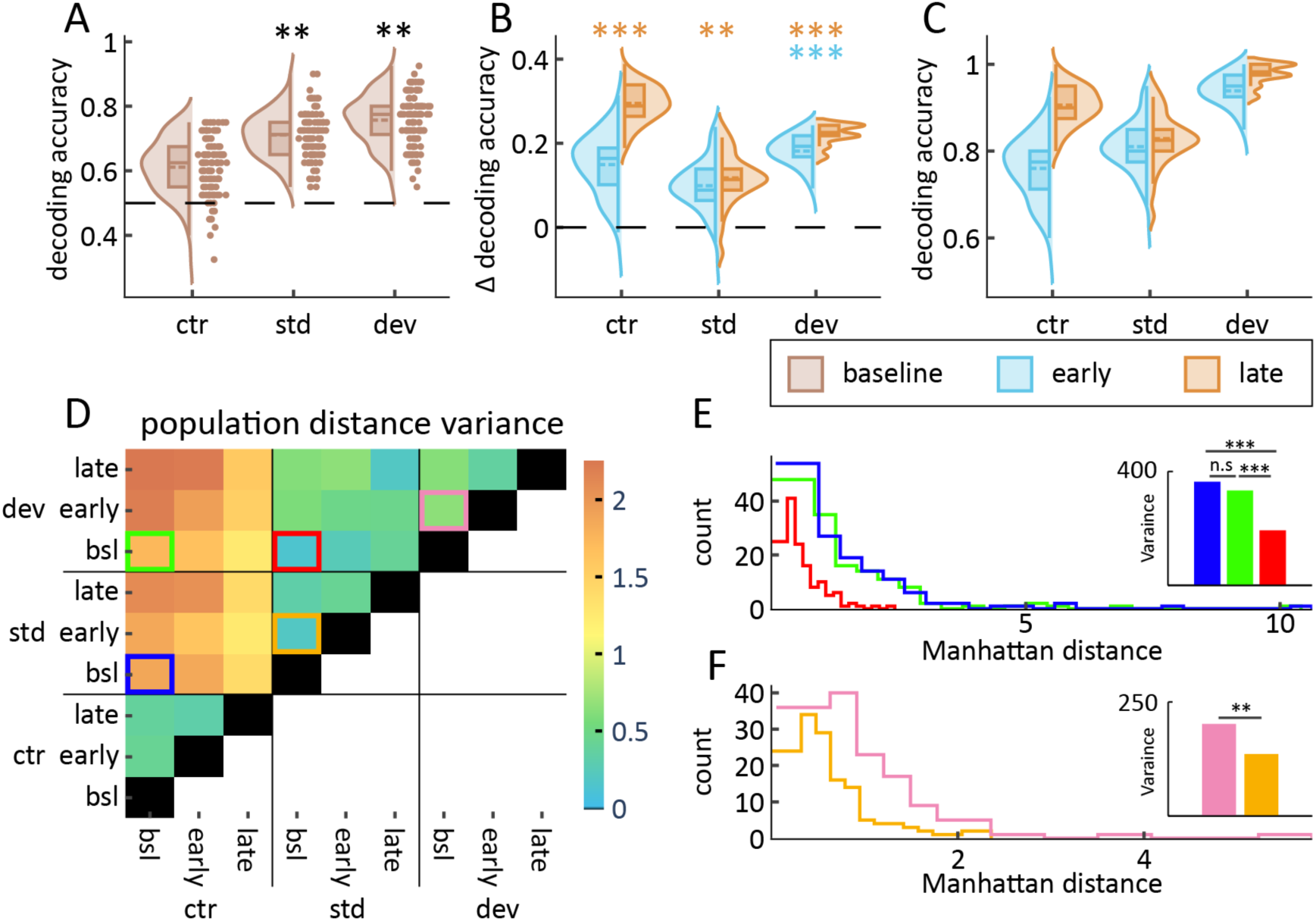
Regularity and mismatch have opposing effects on population stability while improving stimulus-specific population responses. (A) Orientation decoding accuracy at baseline (−500-0 ms). Each datapoint shows one out of 100 iterations during which decoding accuracy was estimated. The dashed line indicates chance performance. (B) Change in decoding accuracy, relative to trial-type specific average at baseline, during early (left-facing) and late (right-facing) epochs. (C) Absolute orientation decoding accuracy at early (left-facing) and late (right-facing) epochs; no asterisks are shown, as significance was assessed in panels A and B. (D) Variance of the Manhattan distance distribution for all combinations of conditions. Colored boxes indicate the contrast statistically tested and displayed in panels E and F (see Supplementary Table 19). Orange and pink compare baseline to early activity within the std and dev trial types. (E) Manhattan distance distributions of all trial type contrasts at baseline. Colors correspond to those in D; asterisks indicate significant variance differences for the corresponding contrasts. (F) Manhattan distance distributions for baseline-to-early contrasts in the standard (orange) and deviant (pink) trial types; asterisk indicate significant variance differences. *p < 0.05, **p < 0.01, ***p < 0.001; asterisks in panels A-C indicate significance from chance (permutation test, see Methods and Supplementary Table 18); asterisks in panel insets (E-F) indicate significant variance differences (median-based Levene’s test, see Methods and Supplementary Table 19).

Next, we assessed the dissimilarity of population activity between trial types and epochs. We computed dissimilarity using a Manhattan distance metric^56^ (Figure S2). Briefly, for each neuron, we calculated Manhattan distance by selecting two conditions, computing the difference in firing rates for horizontal and vertical gratings (these gratings were presented in all std, dev, and ctr conditions; see *Methods*), and then summing these differences. Within each condition, firing rates were z-scored to allow direct comparison across conditions. Thus, each neuron’s Manhattan distance reflects the sum of the changes in the z-scored activity evoked by horizontal and vertical stimuli. We computed this metric for all cells, resulting in a distribution of distances whose variance served as a measure of population-level dissimilarity.

During baseline, we found that the variance of population dissimilarity between standard and deviant conditions (the MMN contrast) was lower than for either the control-standard (SSA) or deviant-control (DD) contrasts (Figure 3E, inset; Supplementary Table 19, *p* < 0.001 from MMN vs. SSA and MMN vs. DD). This indicates that baseline activity patterns were more similar between standard and deviant conditions, supporting the idea that expected stimuli evoke more convergent population states. In contrast, both control-standard and deviant-control comparisons showed greater variance, indicating more heterogeneous neural responses when expectation was not established.

Further, among both control and oddball trial types, we observed a subset of neurons exhibiting firing rate shifts to vertical and horizontal stimuli exceeding four standard deviations. These results indicate that repeated sequences of standard stimuli constrained population activity in a stimulus-specific manner. Together with the decoder results, they suggest that before stimulus onset, repeated exposure to the standard sequence shaped V1 activity into a more stimulus history-constrained pattern, yet retains orientation-specific information from prior presentations.

When comparing baseline to the early response epoch, we found that Manhattan distance was significantly lower for standard than for deviant conditions (Figure 3F, inset; Supplementary Table 19, *p* = 0.0078). This indicates that deviant stimuli, which violate expectations, induce more substantial and variable shifts in orientation-tuned population activity than standard stimuli, which align with prior input history. Further analysis of the distance distributions revealed that this effect was partly driven by a subset of neurons exhibiting exceptionally large shifts in firing rates.

### V1 Excitatory Neurons Exhibit Trial Type-Dependent Changes in Orientation Preference

The preceding results indicate that population activity patterns in V1 can shift across trial types. However, it remains unclear whether these shifts reflect non-selective changes in firing rates or modifications in stimulus selectivity at the single-neuron level. To further examine this, we first quantified each neuron’s stimulus selectivity using the orientation selectivity index (OSI), a classical metric that summarizes the degree to which a neuron preferentially responds to a particular orientation over others (e.g., 57,58). Here, OSI was computed for all trial types and epochs, allowing us to assess changes in the proportion of highly selective neurons as a function of experimental context. We found that this proportion varied across and showed a positive correlation with the Manhattan distance between the trial-averaged activity patterns across the recorded neurons (Figure S3), suggesting that shifts in population-level responses are associated with corresponding changes in stimulus selectivity.

However, because our oddball task primarily involved repeated presentation of horizontal and vertical gratings, we sought a more targeted measure of orientation tuning for these specific stimulus axes. We therefore computed the orientation preference index (OPI), which quantifies the relative preference for horizontal versus vertical orientations (see *Methods*), for the same population of excitatory neurons used in the decoding analyses (N = 141). For example, a neuron that responds more strongly to horizontal than vertical stimuli would have a positive OPI, while a neuron with stronger responses to vertical gratings would have a negative OPI. This approach enabled us to track context-dependent changes in individual neuron’s orientation preference across trial types—a key focus of our subsequent analyses.

Notably, as illustrated by the example in Figure 4A, orientation preference could switch between trial types; this cell responded preferentially to horizontal gratings under standard and deviant conditions, but preferred vertical gratings in the control condition. A second example, shown in Figure 4B, highlights a neuron that exhibited a preference switch from standard to deviant trials specifically during the late epoch. The significance of each cell’s OPIs was assessed through permutation analysis at *p* < 0.05. The number of neurons with significant OPI increased from early to late epochs, with similar proportional distributions across conditions (Figure 4C; Supplementary Table 20).

**Figure 4.**
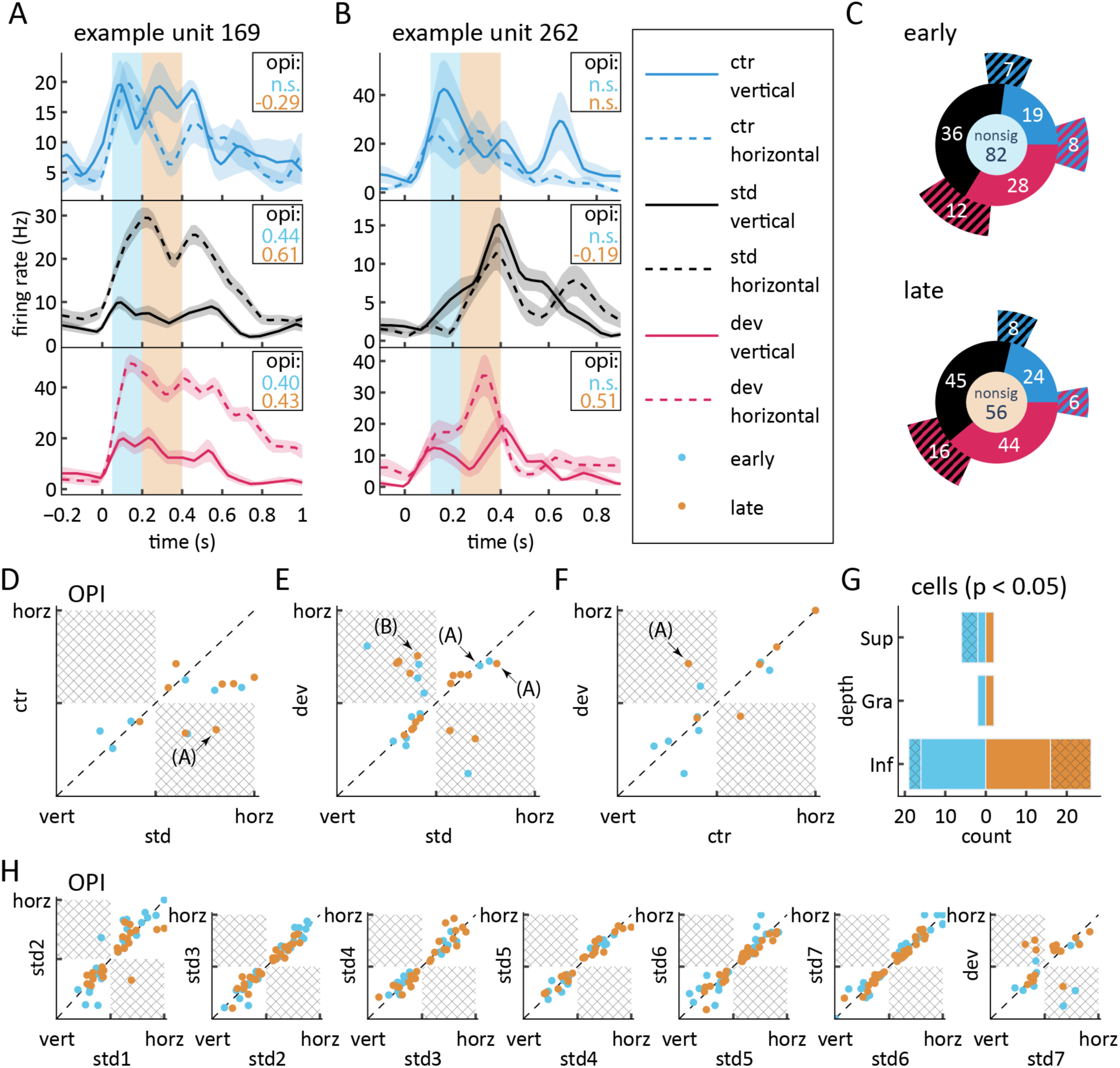
Predictability and mismatch evoke stimulus preference switches in a subset of extragranular V1 neurons. (A and B) Peri-stimulus time histograms (PSTHs, mean ± SEM) for two example neurons exhibiting preference switching. Curves are shown for each stimulus and trial type. Shaded marker areas indicate early and late epochs. Timeseries were smoothed using a 100 ms Tukey window. (C) Sunburst charts showing the distribution of neurons across tuning classes for early (top) and late (bottom) epochs. The center value denotes the number of untuned neurons. Inner ring segments represent numbers of neurons tuned in only one trial type; outer ring segments indicate neurons tuned to in combination of trial types. Striped colors refer to the color code used in (A) (red: deviant; blue: control; black: standard). Neurons represented by the outer ring were analyzed in panels D-G. (D-F) Change in stimulus preference and selectivity strength as a function of trial type comparison. X and Y axes range from perfect vertical preference (−1) to perfect horizontal preference (+1). The diagonal line denotes the null hypothesis: stable preference and tuning strength irrespective of stimulus context. Each point represents a single neuron’s OPI in a pairwise comparison between two trial types. Data points in the highlighted quadrants indicate preference-switching cells (see main text). The asteriks (A) and (B) denote the data points for the example cells shown in panels A and B. Note: Selectivity strength is defined as the absolute value of the OPI, quantifying the degree to which a neuron is tuned toward one orientation axis. (G) Distribution of stable and preference-switching neurons by cortical layer for early and late epochs. Checked bars indicate the number of preference-switching cells;White vertical lines within bars denote the total number of neurons sampled per layer. (H) Control analysis: Change in stimulus preference for consecutive standard presentations (e.g., std7 is the final standard in a sequence). Visual conventions as in panels D-F. Only standards 1-7 are shown. The rightmost panel compares the last standard to the subsequent deviant, illustrating the effect of stimulus mismatch on OPI.

We next examined whether neurons exhibited stimulus context-dependent changes in stimulus preference. To ensure that our analysis focused on clear cases of preference shifts, we included only neurons with significant OPI in two or three trial types, allowing for one or three pairwise comparisons of a neuron’s OPI between trial types, respectively. For both early and late latency windows, the numbers of included comparisons are shown in the outer rings of Figure 4C. Using these samples, we plotted OPI values as a function of trial type and identified 7 cases (26%) of significant preference shifts during early latencies and 10 cases (33%) during late latencies (Figure 4D-F). The proportion of neurons exhibiting preference switches between standard and deviant conditions was significantly greater than expected by chance for both early (χ^2^ = 6.32, *p* = 0.012) and late (χ^2^ = 7.39, *p* = 0.007) epochs. Preference switches also occurred among the limited number of double-tuned neurons in the control-standard and control-deviant contrasts, but these did not reach statistical significance (Supplementary Table 21). The largest proportion of preference switches occurred between the standard and deviant conditions. All neurons with preference switches were located in the extragranular layers, although the small sample size of L4 neurons precludes firm conclusions about the absence of preference-switching cells there. Late preference switches were observed exclusively in infragranular neurons (Figure 4G). Reversals of preference from early to late latency windows within the same condition were infrequent, occurring in just 1.2% of all neurons (χ^2^ = 5.03, *p* = 0.025). This rate was significantly higher than the rate expected under the null hypothesis (see *Methods*), indicating that, while rare, such reversals do occur more often than predicted by chance alone.

To test whether OPI shifts reflected genuine stimulus context effects rather than chance fluctuations, we computed OPI values for the sequence of standard stimuli and found only 1 neuron (0.7%) exhibiting an early and 3 neurons (1.5%) exhibiting a late preference switch between consecutive standards (Figure 4H). This specificity to temporal stimulus context suggests that preference switches were evoked by changes in stimulus predictability and mismatch. To further support this interpretation, we compared the observed difference in proportions of OPI inversions among standards trials to those observed between standards and either control or deviant conditions using a permutation analysis. For all comparisons, the observed differences in the proportion of neurons exhibiting preference switches between conditions fell outside the 95% confidence intervals generated by permutation (Figure S4; early control-to-standard: CI = −0.029 to 0.120, observed = 0.15; early deviant-to-standard: CI = −0.027 to 0.118, observed = 0.178; late control-to-standard: CI = −0.066 to 0.167, observed=0.136; late deviant-to-standard: CI = −0.045 to 0.152, observed = 0.172). Finally, to quantify changes in tuning selectivity regardless of the neurons’s preferred direction, we compared the OPI across trial types. Selectivity increased only in the late deviant compared to the late standard condition (Wilcoxon’s matched-pairs signed rank test, *p* = 0.026), highlighting that, in addition to preference switches, V1 neurons exhibit a sharpening of orientation tuning specifically in response to unexpected stimuli during late processing stages.

To summarize, our findings reveal that orientation preference in V1 neurons is not a static property, but can dynamically switch in response to changes in stimulus regularity and predictability, with the greatest flexibility observed between standard and deviant contexts. Notably, this dynamic remapping of tuning occurs alongside sharpening selectivity specifically during late deviant trials, suggesting that the V1 network is capable of adaptively enhancing both the strength and specificity of its sensory representations when faced with unexpected changes in the environment. These results extend classical views of sensory tuning by showing that, even in primary visual cortex, neuronal feature preference and selectivity are dynamically shaped by temporal context and sensory predictability. To further explore the functional consequences of this tuning flexibility, we next examined its impact on population-level stimulus encoding.

### Stable and Preference-Switching Cells Encode Similar Amounts of Task-Relevant Information

To assess the contribution of preference-switching cells to stimulus orientation encoding, we split the single-neuron dataset into two complementary subsets: one containing preference-switching neurons and the other containing matched stable-preference neurons. Stabled neurons were matched to preference-switching neurons based on similar firing rates and trial counts. We then compared decoding performance between these two groups. Across all combinations of trial type and epoch, decoder accuracy was similar between data splits containing preference-switching neurons and their matched stable controls (Figure 5A-F; Supplementary Tables 22-23), with no significant differences observed (all *p* > 0.05).

**Figure 5.**
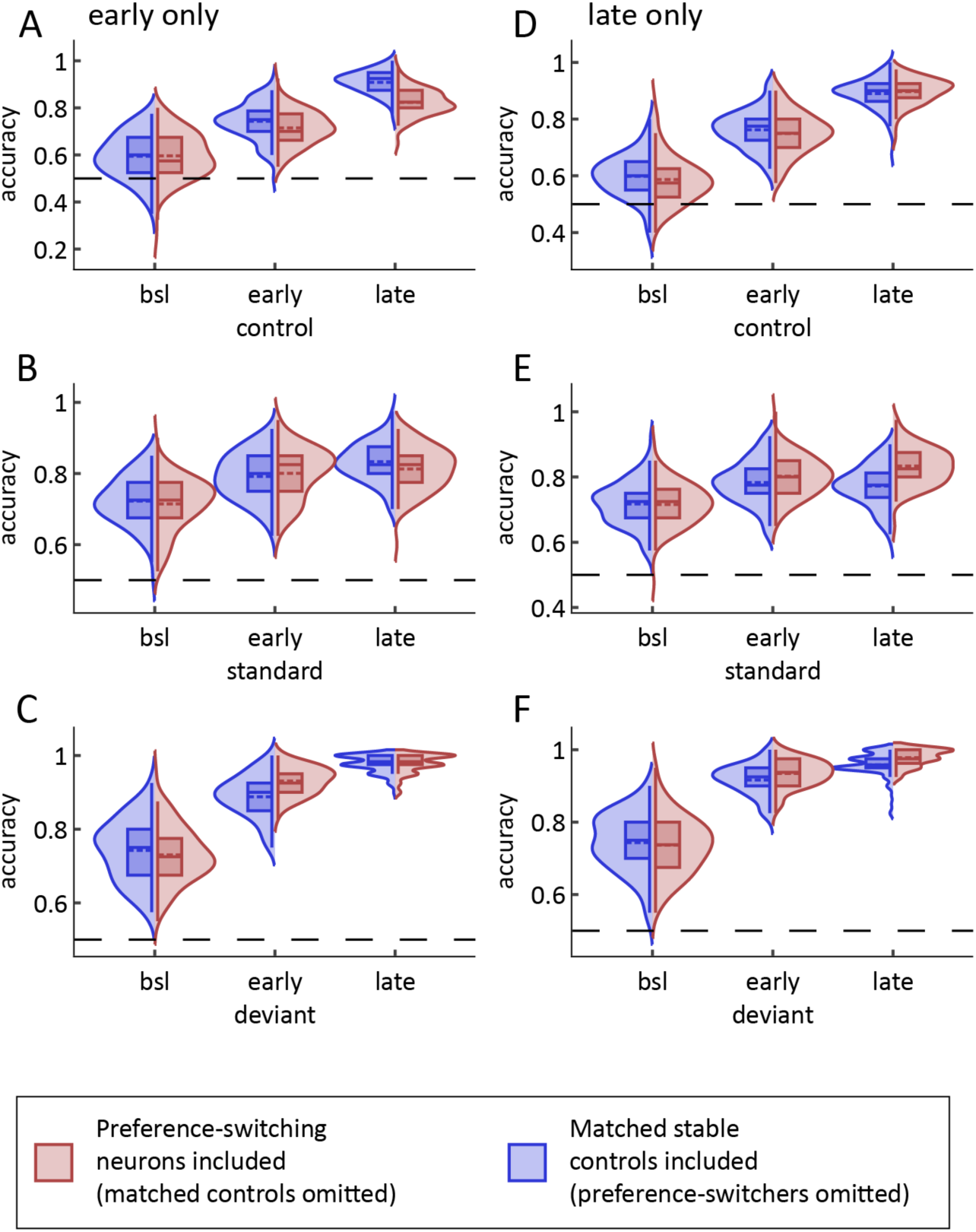
Preference-switching cells do not improve or worsen stimulus decoding performance. (A-F) Comparison of orientation decoding accuracy between complementary dataset splits at baseline (bsl), early, and late epochs. The horizontal dashed line indicates chance preformance. (A-C) Decoding performance for control (A), standard (B), and deviant (C) trial types during the early epoch, with either preference-switching cells included (matched controls omitted, blue) or matched stable controls included (preference-switchers omitted, red). (D-F) Same as A-C, but for the late epoch. Performance was assessed using k-nearest neighbors classification of orientation (horizontal vs. vertical) based on single-neuron activity. No significant differences were found between data splits (all p > 0.05; see Supplementary Tables 22-23).

To independently validate these findings, we also performed a control analysis focusing exclusively on preference-switching and their matched stable counterparts. As expected, restricting the analysis to these smaller subsets led to lower overall decoding performance, though mean accuracy consistently exceeded chance. Preference-switching neurons showed slightly lower decoding accuracy in the early epoch, but modestly outperformed matched controls in the late standard and early deviant conditions. However, none of these differences reached statistical significance (Figure S5; Supplementary Tables 24-25).

Together, these analyses demonstrate that preference-switching neurons are just as capable as stable neurons in supporting robust orientation decoding at the population level, even when directly compared or considered in isolation. This indicates that context-dependent shifts in orientation preference do not impair the ability of the V1 population to encode stimulus orientation.

## Discussion

This study aimed to investigate how neuronal stimulus representations in V1 are influenced by stimulus predictability and mismatch, with a focus on laminar and population-level dynamics. Using simultaneous recordings of single-unit activity and LFPs in awake, head-fixed mice, we identified distinct neural signatures of SSA, MMN, and DD. Our results reveal a temporal and spatial dissociation of DD signal across cortical layers. Early DD responses, as reflected in the ERP (LFP), were localized to supragranular layers (L2/3), while late DD responses in single-neuron activity emerged predominantly in infragranular layers (L5/6). In addition, the informational content and population activity patterns varied systematically with stimulus predictability and deviance, in a manner consistent with predictive coding and cortical microcircuit models of deviance detection^4,52^. Notably, we report a novel phenomenon in which V1 neurons shift their orientation preference depending on the temporal context of stimulus presentation. Such context-sensitive preference shifts may serve to enhance V1’s capacity to encode unexpected events, supporting flexible and dynamic sensory representations.

Such context-driven preference switches may function to amplify weak or ambiguous inputs during mismatch, allowing the system to signal deviations from predicted features more robustly. Mechanistically, this flexibility could arise either from dendritic integration within V1 pyramidal cells or through modulatory feedback that alters thalamic processing. For example, corticothalamic projections from V1 to the LGN may influence how incoming visual information is gated and temporally structured, thereby shaping the input that ultimately reaches cortex^59,60^.

### Predictive coding and deviance detection

PC offers a framework of cortical information processing that aligns with findings from oddball paradigms^7,9,10,22,23,26^. It has guided recent efforts to identify the neuronal mechanisms underlying DD^1,3,26,31,32,46^. A central tenet of PC is that non-local feedback inputs to sensory cortex convey predictions about upcoming stimuli^33^. Supporting this, animal studies using visual and visuo-motor mismatch tasks have identified the anterior cingulate cortex (ACC)^26–28^ and the pulvinar^31^ as key sources of feedback to V1. Notably, top-down signals from ACC to V1 emerge with task experience^28,61,62^ and correlate with predicted visual input^27,28^.

In a particularly compelling study, Furutachi et al.^31^ showed that silencing pulvinar projections to vasoactive intestinal peptide (VIP) interneurons in V1 disrupted the enhanced response to deviant stimuli, providing strong evidence that predictive feedback is necessary for DD. Moreover, somatostatin-positive (SST) interneurons have been shown to modulate feedback processing by inhibiting parvalbumin-positive (PV) interneurons, which in turn target pyramidal-cells^25^. Altogether, these findings provide converging evidence that feedback pathways modulate deviance detection in V1, consistent with predictive coding accounts of sensory processing.

Our population-level findings are compatible with the above studies and support the idea that stimulus regularity elicits repeatable, stimulus-specific neural patterns shaped by prior expectations. Notably, in standard and deviant trials—but not in control trials—we successfully decoded the upcoming stimulus from baseline activity in the putative excitatory V1 neuronal population. While these finding is consistent with previous studies for standard and control conditions^25–27,38^, decoding accuracy in the deviant condition may seem counterintuitive and therefore warrants further explanation. Typically, the greater the number of possible deviants, the harder it is to predict which one will appear. However, in our case, there were only two possible stimuli. In this context, decoding accuracy in the deviant (DD) condition likely reflects the decoder’s ability to infer the only possible alternative stimulus based on the preceding regular pattern. That is, for the decoder, the standard and deviant conditions were computationally equivalent: rather than learning a prediction per se (as in the standard condition), it could correctly classify the deviant by identifying the only remaining option in a two-item sequence. This interpretation is supported by the dissimilarity analysis (Figure 3), which showed that baseline population activity for standards and deviants was more similar to each other than to the control condition. One might argue that comparisons between the oddball and many-standards tasks could be confounded by representational drift^63^, given that they were run sequentially. However, recordings for both tasks were conducted within thirty minutes of each other, and although representational drift can occur on this timescale, its effects on responses to simple stimuli are known to be relatively minor^64,65^. Altogether, our results show that temporal regularity in the stimulus train generated non-evoked and stimulus-specific population activity at baseline. After approximately six consecutive standard trials, the deviant stimulus also became increasingly predictable, suggesting that expectation-related processes evolved dynamically within the sequence.

A core feature of classic PC, incorporated into circuit-level accounts of DD^1,27,31,32^, is that feedforward and feedback inputs exert opposing effects within local circuits (for alternative frameworks, see 6,33). This mechanism ensures that top-down predictions suppress the expected components of sensory input, effectively ‘explaining them away’, so that only unexpected, deviant features propagate up the cortical hierarchy as prediction errors.

Based on this, we expected two outcomes. First, deviant stimuli elicit stimulus-specific prediction error signals, thereby enhancing decoding accuracy. While previous studies^26,27,31,32^ have reported general increases in neural responses to deviant stimuli, our results provide direct evidence that DD enhances the encoding of stimulus-specific orientation information at early latencies.

Second, the mismatch between top-down predictions and bottom-up sensory input should result in larger changes in population activity in deviant versus standard conditions. Consistent with our first prediction, decoding accuracy at early latencies significantly improved on deviant, but not control or standard, trials (Figure 3B). In line with our second prediction, population orientation coding shifted more strongly from baseline in the deviant condition compared to the standard condition (Figure 3F). Both these population-level effects emerged during the transition from baseline to the early window and were mirrored at the single-neuron level by a positive correlation between Manhattan distance and orientation selectivity (Figure S3).

### Orientation preference switches in a subset of neurons in the primary visual cortex

Surprisingly, we found that in a subset of neurons (26% during early latencies and 33% during late latencies), stimulus regularity and deviance altered which orientation elicited the strongest response compared to baseline. These preference switches, restricted to layers L2/3 and L5/6, involved complete reversals between two orthogonal orientations and were not consistent with classical adaptation effects like the tilt-aftereffect (TAE), which typically induces only minor (∼10–20°) shifts in preferred orientation^66–69^.

The TAE, a well-documented perceptual phenomenon, involves a shift in perceived orientation following exposure to a slightly offset adaptor. At the neural level, it is associated with subtle tuning curve shifts, likely mediated by local adaptation mechanisms^67^. In contrast, the dramatic 90° preference switches observed in our study here suggest a different mechanism, possibly involving context-dependent top-down inputs that transiently reshape stimulus preference via feedback-driven modulation^31^.

While prior studies, such as those of Keller and colleagues^29,32^, distinguished between neurons encoding predicted versus actual stimuli, they did not report neurons changing their tuning preferences across conditions. Their work emphasized how L2/3 neurons integrate top-down motor-related and bottom-up visual input but maintained fixed stimulus selectivity. In contrast, our findings show that stimulus selectivity itself can dynamically shift depending on temporal context, with some neurons switching orientation preference between standard and deviant trials. This suggests that rather than representing stable feature detectors, a subset of V1 neurons may adaptively remap their tuning to align with evolving predictions during sensory processing.

These dynamic shifts in orientation preference point to a circuit-level mechanism through which predictive context can modulate sensory tuning properties in V1. While this phenomenon recalls the context-driven representational changes observed in the hippocampus^70^, its presence in early sensory cortex suggests a departure from classical views of fixed tuning during perception. In contrast to classical predictive coding models, where prediction errors arise from mismatches between stable representations and sensory input, our results imply that the representations themselves may be reconfigured based on contextual expectations. Such adaptability could be mediated by feedback inputs that selectively modulate synaptic efficacy or local inhibition, allowing certain neurons to realign their tuning in response to contextual violations. Notably, despite their shifting preferences, these neurons retained high stimulus selectivity, indicating that tuning flexibility does not come at the cost of information content but may instead serve to optimize it under dynamic sensory contingencies.

The present results establish that preference-switching neurons contribute reliable stimulus-specific encoding across conditions, suggesting that their tuning flexibility serves a functional role in adapting to contextual changes. To clarify the mechanisms and conditions under which such remapping occurs, future studies should employ a broader set of orientations and directly manipulate top-down influences.

### Laminar Differences

We partitioned responses into different cortical layers and temporal stages. In line with other studies, we observed early DD in supragranular LFP, underscoring the role of L2/3 in reconciling expected versus actual inputs^4,27,29,31–33,46^. DD at this early latency suggests that it could be locally generated or depend partly on spontaneous activity already present at response onset.

We also identified late responses in infragranular layers that, to our knowledge, have not been reported previously. Specifically, late infragranular DD was observed in single-unit activity but not in the ERP signal, with 10 of the 26 affected neurons exhibiting orientation preference switches across trial types. The discrepancy between the presence of DD in single-units and its absence in LFPs likely reflects the distinct physiological processes captured by these signals: LFPs primarily reflect synaptic input and dendritic processing, whereas SUA represents the spiking output of individual neurons. This suggests that infragranular neurons may receive similar input across trial types, yet differ in their output under conditions of deviance. Comparing infragranular DD findings across studies remains challenging, given variations in task design, recording techniques, and anatomical targeting.

Previous studies have generally reported fewer or weaker DD effects in infragranular layers, particularly in visual cortex^25,26,29,32^. For instance, some have found no trial type-dependent differences in CSD or MUA responses at infragranular depths, and only a small proportion of L5/6 neurons showed DD in calcium imaging, compared to supragranular layers. Our findings expand on this by revealing late DD responses in infragranular single-unit activity, despite the absence of corresponding effects in ERP signals. This discrepancy underscores the potential for distinct circuit dynamics in deep layers, which may not be readily captured by synaptic-level signals. Interestingly, across studies using LFP or calcium imaging, control stimuli often evoke stronger responses with increasing cortical depth^32^ —a pattern we also observed. Whether this depth-dependent sensitivity reflects specialized processing features of infragranular layers processing or specific computational effects of the many-standards control remains an open question. Future studies employing alternative oddball designs may help clarify the circuit mechanisms underlying these late infragranular DD effects.

### Functional and Methodological Implications of Preference Shifts

The presence of preference-switching cells raises questions about how to define a neuron’s orientation selectivity. If a cell prefers stimulus X in the standard condition but switches to stimulus Y in the deviant condition, which preference should be considered canonical? Our use of the many-standards control task helps to clarify this issue. Because stimuli in this condition are presented in random order and are not embedded in a predictable sequence, it minimizes the influence of predictive processes on neuronal responses. This allows the many-standards condition to serve as a neutral reference for functional characterization, similar to classical paradigms that assess tuning using pseudorandom, equiprobable stimulus presentations. These preference shifts may reflect context-sensitive coding rather than instability per se.

From a computational perspective, such cells could flexibly encode internal representations or carry prediction-related signals, rather than the stimulus in a fixed manner. While their precise function remains to be clarified, our findings challenge the traditional view of static feature selectivity in V1 and suggest that orientation coding may be dynamically reshaped by contextual demand, revealing a greater degree of representational flexibility than previously assumed.

The existence of preference-switching cells challenges traditional methods of computing SSA, MMN, and DD for single-cell data. These measures are typically derived by contrasting trials with identical stimuli, which, in counterbalanced designs, yields separate contrast effects for each stimulus type. Current approaches to summarize these effects, such as averaging across stimulus types or selecting the stimulus that produces the largest effect, are appropriate for population-level measures like EEG, MEG. However, in single-neuron analysis, these methods risk conflating stimulus tuning properties with the effects of predictability and deviance. This issue, first highlighted by Nelken and Ulanovsky^11,12^, is further underscored by our finding that, for some neurons, stimulus preference is dependent on temporal stimulus context. The presence of preference-changing cells emphasizes the need for future studies to adopt strategies that account for changes in feature coding, thus providing a more accurate understanding of the neural mechanisms underlying oddball responses.

### Conclusions

Predictability and prediction error profoundly shape how stimuli are processed, influencing the neuronal circuits that underlie these effects. Here, we report temporally and spatially specific signatures of SSA, MMN, and DD across cortical layers in V1. Notably, the emergence of deviance detection in late infragranular responses suggests that deeper layers contribute to the processing of unexpected sensory input. Furthermore, stimulus deviance was associated with enhanced orientation decoding and distinct shifts in population activity patterns. These findings are compatible with both computational models of PC^2,6,8,71^ and broader circuit-level frameworks of deviant detection^1,32,51^. Together, they suggest that predictable contexts give raise to consistent stimulus-specific neural activity patterns that interact with incoming sensory input already at early response latencies.

Finally, we observed distinct latency- and layer-specific effects of stimulus context on orientation preference, suggesting that V1 feature coding can shift dynamically depending on predictability and deviance. Such dynamic tuning could arise from the integration of feedforward and feedback inputs across the dendritic compartments of pyramidal neurons. For example, orientation-selective input arriving in layer 4 and propagating to L2/3 may influence the apical dendrites of L5/6 neurons, whose output is further modulated by contextual feedback targeting their basal dendrites—a configuration that supports context-sensitive modulation of sensory processing^72,73^.

These findings point to the need for mechanistic studies that link such effects to local microcircuit connectivity and the interplay between feedforward and feedback processes. Our results underscore the value of examining all cortical layers with high temporal precision to fully capture the dynamics of deviance detection. Future studies should aim to integrate data across layers, latencies, and experimental paradigms to clarify how predictive coding mechanisms structure cortical responses to unexpected sensory events.

## Methods

### Mice and Surgeries

All experimental procedures were approved by the Dutch Commission for Animal Experiments (*Centrale Commissie Dierproeven*) and by the Animal Welfare Body of the University of Amsterdam (permit number: *AVD1110020172385*). A total of eight male mice (postnatal day [PD] 29-61) were included in this study, covering late juvenile to young adult stages. Mice were bred at the Animal Facility of the University of Amsterdam, socially housed in small groups of siblings, and kept on a 12 h reversed day/night cycle in a temperature- and humidity-controlled room. All experiments were performed during the dark cycle. Mice underwent cranial implantation of a custom-made titanium head bar for head fixation and a small craniotomy over the V1 area. For all surgical procedures, mice were anesthetized using a mixture of isoflurane (250 ml, Raman and Weil, Maharashtra, India) and oxygen with a 3% concentration for induction and 1.5% for maintenance. Buprenorphine (25 µg/kg, SC, Temgesic) was used as intraoperative analgesia. Head bars were installed by removing the scalp hair with a razor and applying local anesthetic and antiseptic to the scalp. After a skin incision, the scalp was prepared and glued to adhere the head bar, which was then secured to the skull with acrylic dental cement (C&B Super-Bond, Sun Medical, Shiga, Japan). Next, a 3 mm craniotomy was drilled over area V1, taking care not to damage the dura, and sealed using silicon elastomer (Kwik-Cast, World Precision Instruments, Sarasota, FL, USA). After both procedures, mice were given one week to recover with food and water ad libitum.

### Intrinsic Optical Imaging

To localize area V1, we performed intrinsic optical imaging (IOI) under lightly anesthetized conditions (0.7-1.2% isoflurane) immediately after head bar implantation^74^. A vasculature image was taken prior to the imaging session, and the cortex was illuminated with monochromatic 630 nm light. Images were acquired at 1 Hz using an Adimec 1000m CCD camera (1004 x 1004 pixels) connected to a frame grabber (Imager 3001, Optical Imaging Inc, Germantown, NY, USA), focused about 500-600 μm below the pial surface. Full-field drifting grating stimuli (0.05 cpd spatial frequency, 1.5 Hz temporal frequency) were presented at eight different orientations for one second. Hemodynamic responses were measured after 8 seconds of baseline, with frames baseline-subtracted, averaged, and thresholded to create a visual cortical map. V1 was marked on the skull using the functional map aligned with the vasculature image. Following IOI, a small craniotomy was performed over functionally defined V1. The recording chamber was then sealed with Kwik-Cast silicon elastomer, and mice were allowed a recovery period of several days before being habituated to head restraint for subsequent experimental sessions.

### Experimental Design

Experiments were performed in a custom-made setup consisting of a metal platform for head fixation and a Polyvinyl chloride (PVC) tube for body stability in front of a LCD monitor (1908FP, Dell, 60 Hz framerate) situated at 25 cm distance from the head^75^. Mice were habituated to this type of head-fixation prior to recordings by daily progressive incremental time spent in head-fixation. Visual stimulation was generated using the Psychophysics Toolbox^76^ in MATLAB 9.13.0 R2022b (The MathWorks Inc, Natick, MA, USA) and presented using *Octave* on a Linux machine. The stimuli consisted of full-field, 50% contrast sine-wave moving gratings (temporal frequency: 2 cycles/second, spatial frequency: 0.05 cycles/degree) presented at two possible orientations (0° and 90°) for 0.5 s, interspersed with a 1.5 s isoluminant gray screen. The oddball task consisted of a repeating pattern of 6-10 consecutive presentations of one stimulus (standard), followed by an orthogonal (deviant) stimulus. After 20 iterations of this pattern, the roles of standard and deviant were swapped. The first two presentations following each swap were excluded from analysis. A *Many-standards* session, consisting of a pseudorandom sequence of moving gratings at eight different orientations (0°, 22.5°, 45°, 67.5°, 90°, 122.5°, 135°, 157.5°) was presented as a control sequence. Each stimulus was presented 20 times. Trials were excluded if their orientation was identical to the preceding stimulus.

### *In vivo* Electrophysiology

Electrophysiological recordings were done in a sound-attenuated and electromagnetically shielded chamber. During recording sessions, full-laminar microelectrode arrays containing 32 electrodes spaced at 100 µm (CM32-A2×16-10mm-100-500-177, Neuronexus, Ann Arbor, MI) were dipped in DiI and gently lowered using a micromanipulator until the entire array penetrated the cortex and the tissue below it (∼1600 µm). The exposed cortex and skull were covered with 1.3-1.5% agarose in artificial CSF to prevent drying and to help maintain mechanical stability during the recording. The ground electrode was connected to the head bar, and the reference electrode contacted the agarose solution. Neurophysiological signals were amplified (x1000), bandpass filtered (0.1 Hz to 9 kHz), and acquired continuously at 32 kHz (hardware: Digital Lynx SX, software: Cheetah5.7.4, Neuralynx, Bozeman, MT). The maximum number of recording sessions was limited to three to minimize recording from damaged tissue.

### Histology

Mice were sacrificed at the end of experiments using an overdose of pentobarbital and transcardially perfused with 4% paraformaldehyde (PFA) in a phosphate-buffered saline (PBS) solution. Brains were sliced into 50 μm coronal sections with a vibratome, stained with DAPI (0.3 μM), and imaged to verify the location of recording sites in the primary visual cortex (Figure 2A).

### Signal Preprocessing

A total of 13 sessions from 8 animals were included in this study (table 26). Two of these recordings yielded no single-unit activity clusters and were omitted. LFPs were extracted from the raw signal by applying a low-pass Butterworth filter at 300 Hz, downsampling to 1 kHz, and applying a notch filter at 50 Hz (including harmonics) to remove line noise. LFP data trials with excessively high or low variance were considered artifactual and discarded. Multi-unit activity (MUA) was obtained by using a second-order Butterworth bandpass filter between 0.5 and 5 kHz, followed by common-average referencing, full-wave rectification, temporal smoothing with a 10 ms Gaussian kernel, and downsampling to 1 kHz. The average current source density profile (CSD) was calculated from the local field potential (LFP) signal using the second spatial derivative, smoothed with a 30 Hz low-pass filter, and spatially interpolated using Modified Akima Cubic Hermite Interpolation^48,49^.

Single-neuron activity was obtained from the raw data using Kilosort 2.0^77^, an automated template-matching spike sorting algorithm. Manual curation of clusters was performed in Phy2^78^. Units were retained if they exhibited consistent waveforms, clear refractory periods, and good separation from noise or multi-unit activity. Neurons were classified into subtypes based on their peak-to-trough duration: units with durations <0.45 ms were labeled narrow-spiking (NS) (putative interneurons), and those >0.55 ms as broad-spiking (BS) (putative excitatory neurons)^45,74^. Units in the intermediate range were left unclassified. Only BS units were included for further analyses. Finally, inclusion in single-neuron analysis required significantly increased firing to at least one stimulus condition relative to baseline (p<.05, one-tailed paired t-test). This criterion was not applied to population-level decoding or tuning analyses.

### Laminar Depth Estimation

Laminar subregions were estimated by sorting channels along their depth and identifying landmarks in the MUA and CSD data. Specifically, the channel with the earliest onset of a MUA response was designated as L4^79^. From the CSD data, the laminar location was determined by aligning the distribution of sinks and sources to pre-established landmarks^50^. We then used these landmarks to estimate the depth of each microelectrode on the probe and overlayed the anatomical borders from the Allen Brain mouse atlas to label each channel as supragranular, granular, or infragranular.

### Calculation of Neural Responses Across Trial Types and Contrasts

Standard (std) and deviant (dev) trials were extracted from the oddball task sequence. Std trials were defined as stimuli immediately preceding a deviant stimulus within the repetitive sequence. Control (ctr) trials were selected from the “many-standards” task sequence, matching the orientation used in the oddball task (i.e., drifting gratings at 0° and 90°). We computed neural responses in three contrast conditions for analysis: Stimulus Specific Adaptation (SSA), Mismatch Negativity (MMN), and Deviant Detection (DD). These contrasts were calculated per unit/channel, trial-, and stimulus-type using the following formulas: SSA = ctr – std; MMN = dev – std; DD = dev – ctr. The resulting trial types and contrasts were averaged across stimulus types, channels/units, sessions, or animals. Trial-averaging and temporal binning or smoothing were applied as mentioned in the results section and figure captions.

### Statistical and Quantitative Analyses Statistical Analyses

Most statistical comparisons were assessed using linear mixed-effects models, which accounted for within-subject variability by including recording session, electrode channel, neuron identity, or layer specificity as random effects. Statistical analyses were conducted in R (version 4.3.0; RStudio 2023.12.1+402 “*Ocean Storm*”) using the *lmerTest* (3.1-33) and *multcomp* (1.4-23) packages. Models were fit using restricted maximum likelihood (REML). Trial type was included as a fixed effect, while session, channel, or neuron were modeled as random intercepts depending on the analysis level (LFP or SUA). Post-hoc comparisons between trial types were adjusted using Holm’s method.

Statistical power was assessed using the *simr* package (version 1.0.7) by assuming fixed effects at 80% of their observed magnitude. Models with at least 80% power^80^ were retained. All model coefficients, test statistics, and adjusted p-values are reported in Supplementary Tables 1-17. When effects are described in the *Results* section, the corresponding tables are cited.

### Other Quantitative Analyses

#### Stimulus Decoding

We employed a k-nearest neighbors (KNN) classifier to decode grating orientation (horizontal vs. vertical) based on binned spike counts (100 ms bins). Baseline activity was defined as the 500 ms before stimulus onset, with early and late epochs corresponding to the first and second 200 ms post-stimulus, respectively. Decoding was performed separately for each trial type and time window using 5-fold cross-validation (80% training, 20% test). Accuracy was determined via majority voting among the five nearest neighbors, using Euclidean distance. This was repeated 100 times to generate a distribution of decoding accuracy. The implementation used Python’s *scikit-learn* package (version 1.3.0). Statistical significance for decoding accuracy was assessed using a resampling-based approach. For each condition, the observed mean accuracy was compared against a null hypothesis value (e.g., 0.5 for chance-level classification or 0 for no change). This comparison used 100 bootstrap resamples to generate a sampling distribution, from which empirical p-values and 95% confidence intervals were computed. (Figure 3A-B) Effect sizes were expressed as bias-corrected standardized mean differences (SMDs).

#### Manhattan Distance and Population Dissimilarity

We calculated Manhattan distance to assess population dissimilarity across conditions^56^. Firing rates were z-scored across neurons for normalization. Each neuron’s distance was computed as the sum of the absolute differences in normalized responses to horizontal and vertical gratings between two trial types (Figure S2):

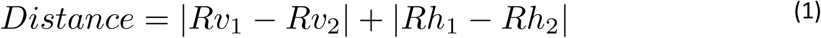

Where *Rv* and *Rh* denote responses to vertical and horizontal stimuli. The resulting distances followed a folded normal distribution. Group differences were tested using a median-based Levene’s test.

A significant *p*-value in Levene’s test indicates a meaningful difference in the variability of neural population responses between contrasts. Here, lower variance in standard versus deviant conditions reflects more consistent (less variable) population activity patterns, suggesting that expectation reduces neural response heterogeneity, while violation of expectation (deviant) increases it.

#### Stimulus selectivity

Orientation preference index (OPI) was computed per neuron and trial type as:

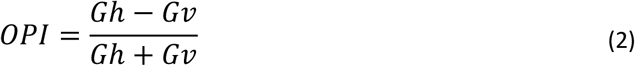

Where *Gh* and *Gv* are z-scored firing rates to horizontal and vertical gratings. Significance was tested using t-tests against null distributions generated from 1000 permutations of stimulus labels. P-values were Bonferroni-corrected for multiple comparisons across neurons and conditions (α = 0.05).

#### OPI Inversion Rate Permutation Test

To assess whether stimulus context altered orientation preference, we compared Orientation Preference Index (OPI) sign changes (i.e., preference inversions from horizontal to vertical or viceversa) between trial types. Analyses included only neurons with significant OPI in both conditions being compared. The inversion rate was defined as the proportion of such doubly significant neurons that change preference. As baseline, we computed the inversion rate for successive standards-to-standards trials. Deviant-standard and control-standard inversion rates were compared against this baseline using a permutation test. In each of 1000 iterations, we shufled conditions and OPIs within neurons and drew random samples with replacement to match the observed sample size. Confidence intervals were Bonferroni-corrected (α = 0.05), and significance was assessed by comparing the observed inversion difference to the permuted distribution.

#### Matched Omission Decoder Analysis

To determine the contribution of preference-switching cells to stimulus orientation encoding, we classified neurons (N = 141) based on the orientation preference index (OPI), assessed via permutation testing (α = 0.05). Neurons were categorized as non-significant (no significant OPI), significant in a single trial type, stable (significant in two or more trial types without preference inversion), or preference-switching (significant in two or more trial types with opposite-sign OPI). Each preference-switching neuron was matched to a stable neuron with a similar firing rate and trial count (“matched stable control”).

We created two complementary data splits: one excluding preference-switching neurons (retaining only stable controls), and one excluding the matched stable controls (retaining only preference-switchers). K-nearest neighbors (KNN) decoding (see above) was performed on each split to predict stimulus orientation (horizontal vs. vertical) for each trial and epoch. Decoding performance was compared using the standardized mean difference (SMD) and permutation-based significance testing.

## Data and code availability

The data and code used in this study are available upon request.

## Acknowledgments

The authors would like to thank Lucas Lumeij and Philip Oosterholt for their support during the experiments, as well as the Technology Center of the University of Amsterdam for their assistance in developing the recording set-up. We are also grateful to László Négyessy, Zoltán Somogiváry, and Luc Gentet for their initial feedback on the project. This study was supported by FLAG-ERA, co-financed by The Netherlands Organization for Scientific Research (Joint Transnational Calls 2015, 2019. Projects: CANON, DOMINO), and by The Netherlands Organization for Health Research and Development (Joint Transnational Call 2023. Project: MONAD) to C.A.B. and U.O., and by the European Union’s Horizon 2020 Framework Program for Research and Innovation (Specific Grant Agreement 785907—Human Brain Project SGA2— to C.A.B. and U.O.).

