## Supplementary Tables X for "Switches in Orientation Coding by Mouse Primary Visual Cortex Neurons Depend on Stimulus Predictability"

#### Author affiliation:

#### Supplementary Figures

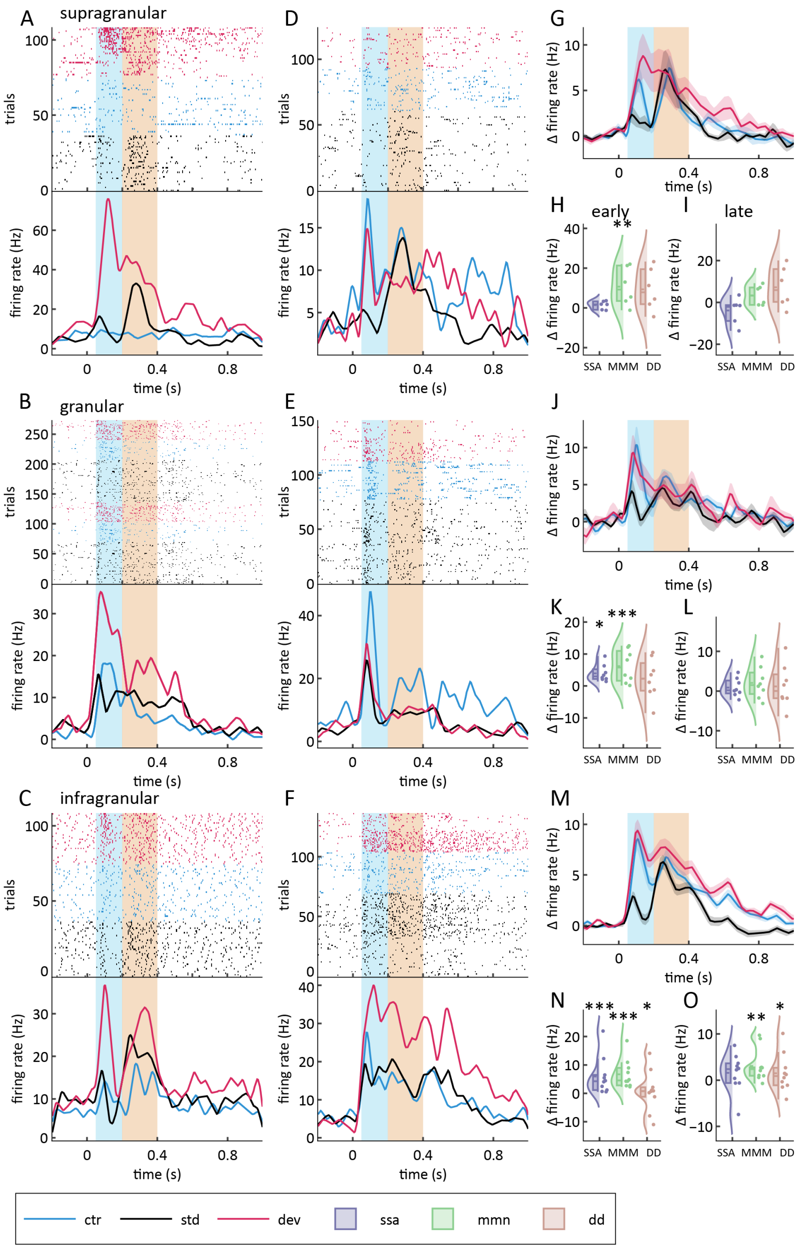


***Figure S1. Laminar spike profile reveals trial type-dependent firing patterns across cortical layers***

*(A–F) Raster plots and peri-stimulus time histograms (PSTHs; mean ± SEM) for six example single units, illustrating trial type-dependent firing responses.*

*(G) Average PSTH of all neurons located in supragranular layers (L2/3) (n = 27).*

*(H, I) Raincloud plots showing the distribution of firing rates across trial types for supragranular neurons during the (H) early and (I) late epochs.*

*(J–L) As in G–I, but for neurons located in granular layers (L4) (n = 18).*

*(M–O) As in G–I, but for neurons located in infragranular layers (L5/6) (n = 76).*

*Raincloud plots are shown in panels H, I, K, L, N, and O. Plotting conventions as in Figure 1.*
*$p < 0.05$, $p < 0.01$, $p < 0.001$ indicate significance from zero (i.e., no change in firing rate), based on linear mixed-effects model analysis (see Methods and Supplementary Tables 12–17).*

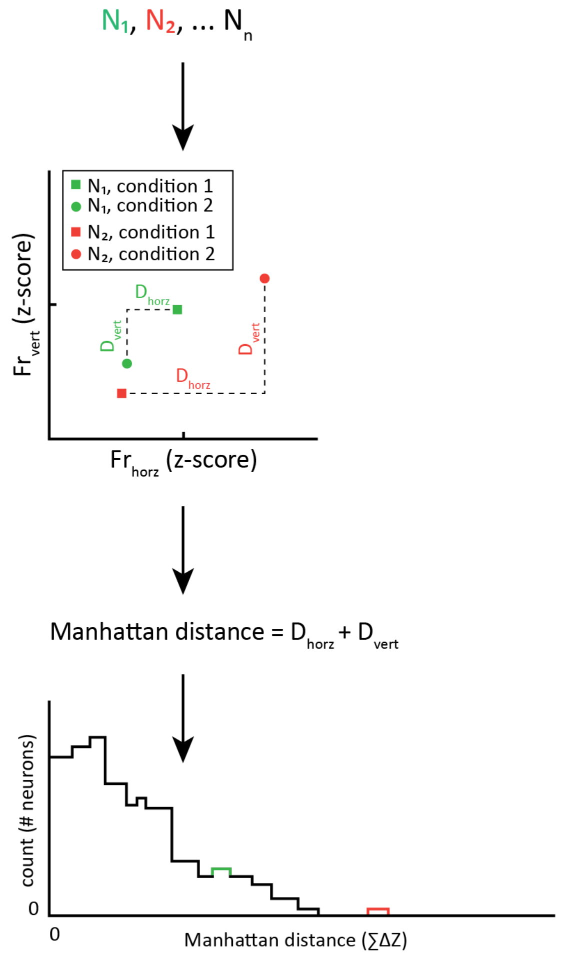


***Figure S2. Schematic of Manhattan distance calculation for population firing rate dissimilarity***

*This diagram illustrates the steps used to quantify population-level dissimilarity between conditions with the Manhattan distance metric.*

*Top: For each neuron ($N_1$, $N_2$, … $N_n$), firing rates (FR) to horizontal (horz) and vertical (vert) gratings were normalized (z-score) separately for each condition.*

*Middle: For each neuron, the Manhattan distance between two conditions was calculated as the sum of the absolute differences in normalized firing rates to horizontal and vertical gratings:*
*Manhattan distance =* $|\Delta FR_{hor}\mathcal{Z}| + |\Delta FR_{vert}|$.

*Bottom: Example histogram illustrating the distribution of Manhattan distance values across neurons for two hypothetical conditions (condition 1: green; condition 2: red). This distribution reflects the degree of population dissimilarity between conditions.*

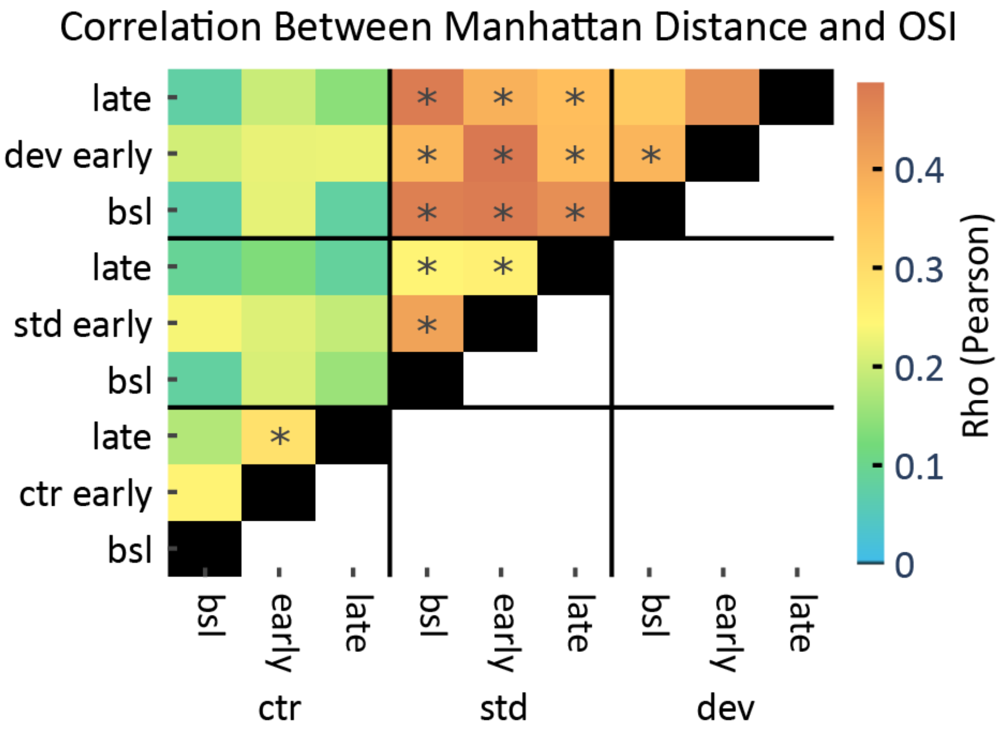


***Figure S3. Population-level dissimilarity correlates with orientation selectivity across conditions***

*Heatmap showing the Pearson correlation coefficient (ρ) between Manhattan distance and orientation selectivity index (OSI) across all pairwise contrasts of trial type and epoch (baseline, early, late) conditions. Each cell displays the correlation for a specific condition pair, with color indicating the strength of the correlation (see color bar).*
*Asterisks indicate statistically significant correlations after Bonferroni correction ($p < 0.05$).*

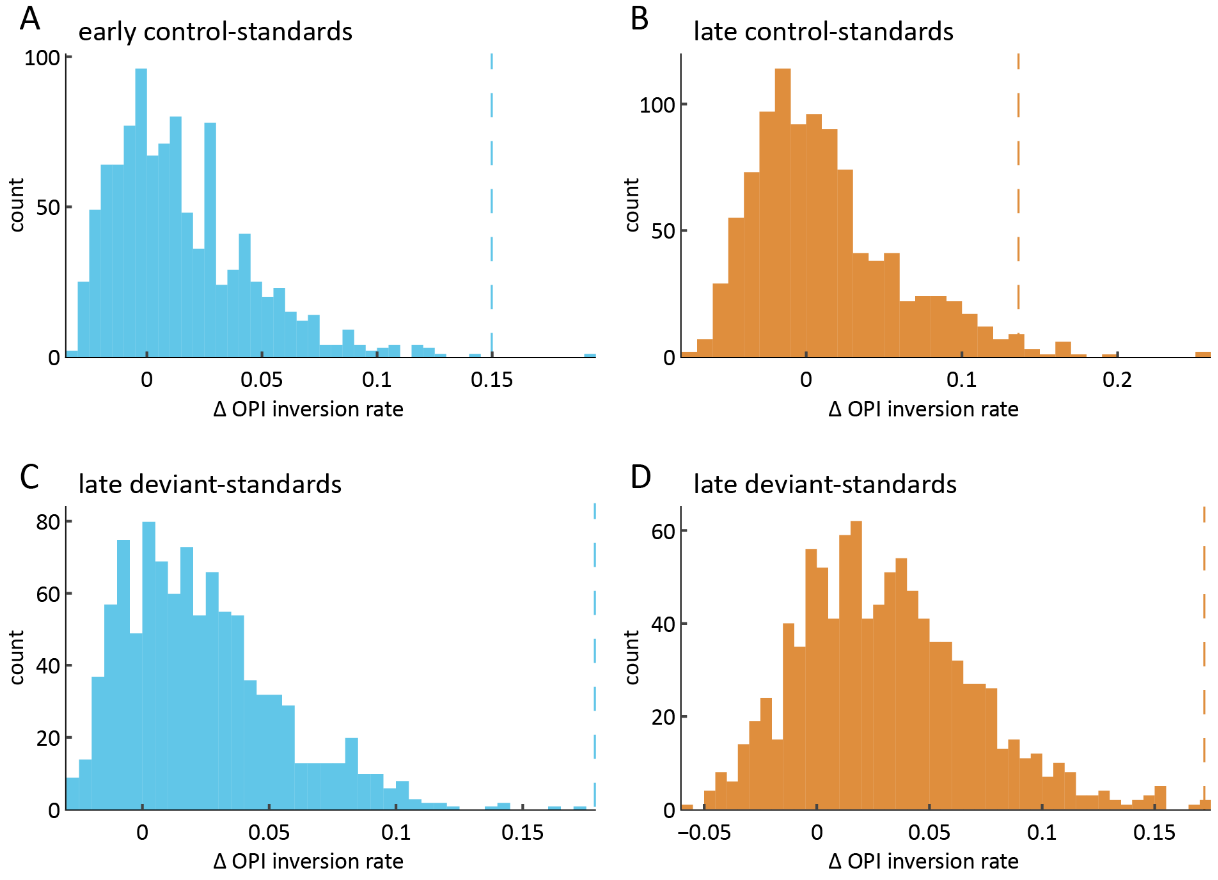


***Figure S4. Permutation analysis supports context dependence of orientation preference switches***

*(A–D) Distributions of permutation-based null values for the difference in OPI inversion rate (Δ OPI inversion rate) between standard and control (A, C) or standard and deviant (B, D) conditions, shown for early (A, B) and late (C, D) epochs.*
*The x-axis (Δ OPI inversion rate) represents the observed minus permuted difference in the proportion of neurons exhibiting preference switches (i.e., OPI inversion) between conditions. Vertical lines mark the observed values.*
*For each comparison, the observed difference fell outside the 95% confidence interval of the null distribution, indicating that preference switches occurred at significantly higher rates in context-changing conditions than expected by chance.*

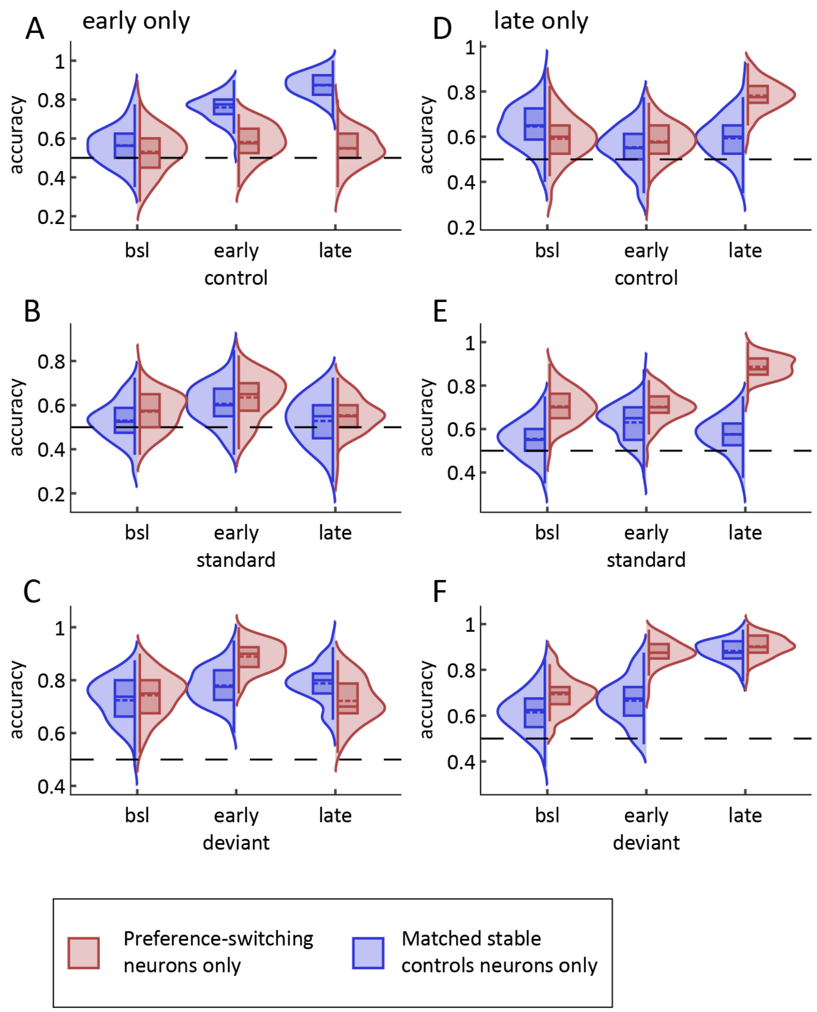


***Figure S5. Decoding performance in populations comprised exclusively of preference-switching or matched stable neurons***

*(A–F) Comparison of orientation decoding accuracy between dataset splits containing only preference-switching neurons (red) or only matched stable controls (blue), at baseline (bsl), early, and late epochs. The horizontal dashed line indicates chance performance.*

*(A–C) Decoding performance for control (A), standard (B), and deviant (C) trial types during the early epoch.*
*(D–F) Same as A–C, but for the late epoch.*

*Performance was assessed using k-nearest neighbors classification of orientation (horizontal vs. vertical) based on single-neuron activity. No significant differences were found between preference-switching and matched stable neuron populations (all $p > 0.05$; see Supplementary Tables 24–25).*

### Supplementary Tables

#### Supplementary Table 1

##### Early ERP Amplitudes for SSA, MMN, and DD Contrasts Across All Channels (Figure 1C)

| Effect | Estimate | Std. Error / SEM | df / z | t/z value | p-value | Adj. p-value (Holm) | Sig. |
|---|---|---|---|---|---|---|---|
| Intercept | -48.22 | 9.01 | 12.36 | -5.35 | 0.00016 | | *** |
| Standard (vs Control) | 25.61 | 1.93 | 502.00 | 13.24 | 1.52e-34 | | *** |
| Deviant (vs Control) | -7.31 | 1.93 | 502.00 | -3.78 | 0.00018 | | *** |
| **Tukey post-hoc contrasts (Holm-corrected p-values)** | | | | | | | |
| Standard – Control | 25.61 | 1.93 | 13.24 | 13.24 | 0 | 0 | *** |
| Deviant – Control | -7.31 | 1.93 | -3.78 | -3.78 | 0.00016 | 0.00016 | *** |
| Deviant – Standard | -32.92 | 1.93 | -17.02 | -17.02 | 0 | 0 | *** |

##### Random Effects of the Linear Mixed-Effects Model

| Group | Name | Variance | Std. Deviation |
|---|---|---|---|
| Channel:Session | Intercept | 417.46 | 20.43 |

Note: Model estimates and standard errors for fixed effects are shown first. Tukey's post-hoc pairwise comparisons (Holm-corrected p-values) are shown below the model estimates. Random effects report variance components for the random intercept. Significance stars: *p < 0.05; **p < 0.01; ***p < 0.001.

#### Supplementary Table 2

##### Late ERP Amplitudes for SSA, MMN, and DD Contrasts Across All Channels (Figure 1D)

| Effect | Estimate | Std. Error / SEM | df / z | t/z value | p-value | Adj. p-value (Holm) | Sig. |
|---|---|---|---|---|---|---|---|
| Intercept | -48.22 | 9.82 | 12.24 | -4.91 | 0.00034 | | *** |
| Standard (vs Control) | 9.96 | 1.71 | 502.00 | 5.81 | 1.08e-08 | | *** |
| Deviant (vs Control) | -0.81 | 1.71 | 502.00 | -0.47 | 0.63540 | | |
| **Tukey post-hoc contrasts (Holm-corrected p-values)** | | | | | | | |
| Standard – Control | 9.96 | 1.71 | 5.81 | 5.81 | 1.22e-08 | 1.22e-08 | *** |
| Deviant – Control | -0.81 | 1.71 | -0.47 | -0.47 | 0.63520 | 0.63520 | |
| Deviant – Standard | -10.78 | 1.71 | -6.29 | -6.29 | 9.58e-10 | 9.58e-10 | *** |

##### Random Effects of the Linear Mixed-Effects Model

| Group | Name | Variance | Std. Deviation |
|---|---|---|---|
| Channel:Session | Intercept | 410.08 | 20.25 |
| Session | Intercept | 1212.20 | 34.82 |
| Residual | NA | 369.92 | 19.23 |

Note: Model estimates and standard errors for fixed effects are shown first. Tukey's post-hoc pairwise comparisons (Holm-corrected p-values) are shown below the model estimates. Random effects report variance components for the random intercept. Significance stars: *p < 0.05; **p < 0.01; ***p < 0.001.

#### Supplementary Table 3

##### Early Firing Rate Differences for SSA, MMN, and DD Contrasts in V1 Single Units (Figure 1G)

| Effect | Estimate | Std. Error / SEM | df / z | t/z value | p-value | Adj. p-value (Holm) | Sig. |
|---|---|---|---|---|---|---|---|
| Intercept | 5.20 | 0.66 | 16.15 | 7.85 | 6.64e-07 | | *** |
| Standard (vs Control) | -3.43 | 0.61 | 240.00 | -5.63 | 4.90e-08 | | *** |
| Deviant (vs Control) | 2.00 | 0.61 | 240.00 | 3.28 | 0.00119 | | ** |
| **Tukey post-hoc contrasts (Holm-corrected p-values)** | | | | | | | |
| Standard – Control | -3.43 | 0.61 | -5.63 | -5.63 | 3.51e-08 | 3.51e-08 | *** |
| Deviant – Control | 2.00 | 0.61 | 3.28 | 3.28 | 0.00104 | 0.00104 | ** |
| Deviant – Standard | 5.42 | 0.61 | 8.91 | 8.91 | 0 | 0 | *** |

##### Random Effects of the Linear Mixed-Effects Model

| Group | Name | Variance | Std. Deviation |
|---|---|---|---|
| Neuron:Session | Intercept | 6.64 | 2.58 |
| Session | Intercept | 1.72 | 1.31 |
| Residual | NA | 22.40 | 4.73 |

Note: Model estimates and standard errors for fixed effects are shown first. Tukey's post-hoc pairwise comparisons (Holm-corrected p-values) are shown below the model estimates. Random effects report variance components for the random intercept. Significance stars: *p < 0.05; **p < 0.01; ***p < 0.001.

#### Supplementary Table 4

##### Late Firing Rate Differences for SSA, MMN, and DD Contrasts in V1 Single Units (Figure 1H)

| Effect | Estimate | Std. Error / SEM | df / z | t/z value | p-value | Adj. p-value (Holm) | Sig. |
|---|---|---|---|---|---|---|---|
| Intercept | 5.42 | 1.06 | 10.02 | 5.11 | 0.00045 | | *** |
| Standard (vs Control) | -0.48 | 0.53 | 240.00 | -0.90 | 0.36800 | | |
| Deviant (vs Control) | 1.25 | 0.53 | 240.00 | 2.35 | 0.01940 | | * |
| **Tukey post-hoc contrasts (Holm-corrected p-values)** | | | | | | | |
| Standard – Control | -0.48 | 0.53 | -0.90 | -0.90 | 0.36700 | 0.36700 | |
| Deviant – Control | 1.25 | 0.53 | 2.35 | 2.35 | 0.03730 | 0.03730 | * |
| Deviant – Standard | 1.73 | 0.53 | 3.25 | 3.25 | 0.00340 | 0.00340 | ** |

##### Random Effects of the Linear Mixed-Effects Model

| Group | Name | Variance | Std. Deviation |
|---|---|---|---|
| Neuron:Session | Intercept | 10.04 | 3.17 |
| Session | Intercept | 8.75 | 2.96 |
| Residual | NA | 17.00 | 4.12 |

Note: Model estimates and standard errors for fixed effects are shown first. Tukey's post-hoc pairwise comparisons (Holm-corrected p-values) are shown below the model estimates. Random effects report variance components for the random intercept. Significance stars: *p < 0.05; **p < 0.01; ***p < 0.001.

#### Supplementary Table 5

##### Control ERP Amplitudes Across Cortical Depth (Figure 2D, G, J)

| Effect | Estimate | Std. Error / SEM | df / z | t/z value | p-value | Adj. p-value (Holm) | Sig. |
|---|---|---|---|---|---|---|---|
| Intercept | -25.43 | 11.28 | 13.08 | -2.26 | 0.04189 | | * |
| Granular (vs Supragranular) | -30.50 | 6.42 | 237.00 | -4.75 | 3.56e-06 | | *** |
| Infragranular (vs Supragranular) | -38.08 | 3.91 | 237.03 | -9.73 | 4.88e-19 | | *** |
| **Tukey post-hoc contrasts (Holm-corrected p-values)** | | | | | | | |
| Granular – Supragranular | -30.50 | 6.42 | -4.75 | -4.75 | 4.11e-06 | 4.11e-06 | *** |
| Infragranular – Supragranular | -38.08 | 3.91 | -9.73 | -9.73 | 0 | 0 | *** |
| Infragranular – Granular | -7.58 | 6.24 | -1.22 | -1.22 | 0.22420 | 0.22420 | |

##### Random Effects of the Linear Mixed-Effects Model

| Group | Name | Variance | Std. Deviation |
|---|---|---|---|
| Session | Intercept | 1538.02 | 39.22 |
| Residual | NA | 843.14 | 29.04 |

Note: Model estimates and standard errors for fixed effects are shown first. Tukey's post-hoc pairwise comparisons (Holm-corrected p-values) are shown below the model estimates. Random effects report variance components for the random intercept. Significance stars: *p < 0.05; **p < 0.01; ***p < 0.001.

#### Supplementary Table 6

##### Early SSA, MMN, and DD Effects in Supragranular Layers (Figure 2E)

| Effect | Estimate | Std. Error / SEM | df / z | t/z value | p-value | Adj. p-value (Holm) | Sig. |
|---|---|---|---|---|---|---|---|
| Intercept | -25.27 | 3.75 | 147.09 | -6.73 | 3.50e-10 | | *** |
| Standard (vs Control) | 13.37 | 2.98 | 190.00 | 4.48 | 1.26e-05 | | *** |
| Deviant (vs Control) | -15.66 | 2.98 | 190.00 | -5.25 | 3.97e-07 | | *** |
| **Tukey post-hoc contrasts (Holm-corrected p-values)** | | | | | | | |
| Standard – Control | 13.37 | 2.98 | 4.48 | 4.48 | 7.31e-06 | 7.31e-06 | *** |
| Deviant – Control | -15.66 | 2.98 | -5.25 | -5.25 | 2.98e-07 | 2.98e-07 | *** |
| Deviant – Standard | -29.03 | 2.98 | -9.74 | -9.74 | 0 | 0 | *** |

##### Random Effects of the Linear Mixed-Effects Model

| Group | Name | Variance | Std. Deviation |
|---|---|---|---|
| Channel | Intercept | 926.50 | 30.44 |
| Residual | NA | 426.63 | 20.66 |

Note: Model estimates and standard errors for fixed effects are shown first. Tukey's post-hoc pairwise comparisons (Holm-corrected p-values) are shown below the model estimates. Random effects report variance components for the random intercept. Significance stars: *p < 0.05; **p < 0.01; ***p < 0.001.

#### Supplementary Table 7

##### Early SSA, MMN, and DD Effects in Granular Layer (Figure 2K)

| Effect | Estimate | Std. Error / SEM | df / z | t/z value | p-value | Adj. p-value (Holm) | Sig. |
|---|---|---|---|---|---|---|---|
| Intercept | -55.93 | 8.64 | 33.95 | -6.47 | 2.13e-07 | | *** |
| Standard (vs Control) | 31.33 | 5.77 | 50.00 | 5.43 | 1.63e-06 | | *** |
| Deviant (vs Control) | -9.65 | 5.77 | 50.00 | -1.67 | 0.10030 | | |
| **Tukey post-hoc contrasts (Holm-corrected p-values)** | | | | | | | |
| Standard – Control | 31.33 | 5.77 | 5.43 | 5.43 | 1.10e-07 | 1.10e-07 | *** |
| Deviant – Control | -9.65 | 5.77 | -1.67 | -1.67 | 0.09410 | 0.09410 | |
| Deviant – Standard | -40.98 | 5.77 | -7.11 | -7.11 | 3.52e-12 | 3.52e-12 | *** |

##### Random Effects of the Linear Mixed-Effects Model

| Group | Name | Variance | Std. Deviation |
|---|---|---|---|
| Channel | Intercept | 1509.61 | 38.85 |
| Residual | NA | 432.13 | 20.79 |

Note: Model estimates and standard errors for fixed effects are shown first. Tukey's post-hoc pairwise comparisons (Holm-corrected p-values) are shown below the model estimates. Random effects report variance components for the random intercept. Significance stars: *p < 0.05; **p < 0.01; ***p < 0.001.

#### Supplementary Table 8

##### Early SSA, MMN, and DD Effects in Infragranular Layers (Figure 2H)

| Effect | Estimate | Std. Error / SEM | df / z | t/z value | p-value | Adj. p-value (Holm) | Sig. |
|---|---|---|---|---|---|---|---|
| Intercept | -63.51 | 3.70 | 185.47 | -17.16 | 3.86e-40 | | *** |
| Standard (vs Control) | 33.51 | 2.68 | 258.00 | 12.48 | 2.71e-28 | | *** |
| Deviant (vs Control) | -0.67 | 2.68 | 258.00 | -0.25 | 0.80440 | | |
| **Tukey post-hoc contrasts (Holm-corrected p-values)** | | | | | | | |
| Standard – Control | 33.51 | 2.68 | 12.48 | 12.48 | 0 | 0 | *** |
| Deviant – Control | -0.67 | 2.68 | -0.25 | -0.25 | 0.80420 | 0.80420 | |
| Deviant – Standard | -34.17 | 2.68 | -12.73 | -12.73 | 0 | 0 | *** |

##### Random Effects of the Linear Mixed-Effects Model

| Group | Name | Variance | Std. Deviation |
|---|---|---|---|
| Channel | Intercept | 1312.80 | 36.23 |
| Residual | NA | 468.27 | 21.64 |

Note: Model estimates and standard errors for fixed effects are shown first. Tukey's post-hoc pairwise comparisons (Holm-corrected p-values) are shown below the model estimates. Random effects report variance components for the random intercept. Significance stars: *p < 0.05; **p < 0.01; ***p < 0.001.

#### Supplementary Table 9

##### Late SSA, MMN, and DD Effects in Supragranular Layers (Figure 2F)

| Effect | Estimate | Std. Error / SEM | df / z | t/z value | p-value | Adj. p-value (Holm) | Sig. |
|---|---|---|---|---|---|---|---|
| Intercept | -32.72 | 3.87 | 134.18 | -8.46 | 4.01e-14 | | *** |
| Standard (vs Control) | 2.08 | 2.74 | 190.00 | 0.76 | 0.44850 | | |
| Deviant (vs Control) | -5.86 | 2.74 | 190.00 | -2.14 | 0.03360 | | * |
| **Tukey post-hoc contrasts (Holm-corrected p-values)** | | | | | | | |
| Standard – Control | 2.08 | 2.74 | 0.76 | 0.76 | 0.44760 | 0.44760 | |
| Deviant – Control | -5.86 | 2.74 | -2.14 | -2.14 | 0.06460 | 0.06460 | |
| Deviant – Standard | -7.93 | 2.74 | -2.90 | -2.90 | 0.01120 | 0.01120 | ** |

##### Random Effects of the Linear Mixed-Effects Model

| Group | Name | Variance | Std. Deviation |
|---|---|---|---|
| Channel | Intercept | 1076.22 | 32.81 |
| Residual | NA | 359.41 | 18.96 |

Note: Model estimates and standard errors for fixed effects are shown first. Tukey's post-hoc pairwise comparisons (Holm-corrected p-values) are shown below the model estimates. Random effects report variance components for the random intercept. Significance stars: *p < 0.05; **p < 0.01; ***p < 0.001.

#### Supplementary Table 10

##### Late SSA, MMN, and DD Effects in Granular Layer (Figure 2I)

| Effect | Estimate | Std. Error / SEM | df / z | t/z value | p-value | Adj. p-value (Holm) | Sig. |
|---|---|---|---|---|---|---|---|
| Intercept | -54.16 | 8.11 | 30.36 | -6.68 | 2.00e-07 | | *** |
| Standard (vs Control) | 5.88 | 4.33 | 50.00 | 1.36 | 0.18070 | | |
| Deviant (vs Control) | 0.15 | 4.33 | 50.00 | 0.03 | 0.97240 | | |
| **Tukey post-hoc contrasts (Holm-corrected p-values)** | | | | | | | |
| Standard – Control | 5.88 | 4.33 | 1.36 | 1.36 | 0.52390 | 0.52390 | |
| Deviant – Control | 0.15 | 4.33 | 0.03 | 0.03 | 0.97230 | 0.97230 | |
| Deviant – Standard | -5.73 | 4.33 | -1.32 | -1.32 | 0.52390 | 0.52390 | |

##### Random Effects of the Linear Mixed-Effects Model

| Group | Name | Variance | Std. Deviation |
|---|---|---|---|
| Channel | Intercept | 1466.03 | 38.29 |
| Residual | NA | 243.63 | 15.61 |

Note: Model estimates and standard errors for fixed effects are shown first. Tukey's post-hoc pairwise comparisons (Holm-corrected p-values) are shown below the model estimates. Random effects report variance components for the random intercept. Significance stars: *p < 0.05; **p < 0.01; ***p < 0.001.

#### Supplementary Table 11

##### Late SSA, MMN, and DD Effects in Infragranular Layers (Figure 2L)

| Effect | Estimate | Std. Error / SEM | df / z | t/z value | p-value | Adj. p-value (Holm) | Sig. |
|---|---|---|---|---|---|---|---|
| Intercept | -58.56 | 4.04 | 165.31 | -14.51 | 2.65e-31 | | *** |
| Standard (vs Control) | 16.60 | 2.43 | 258.00 | 6.84 | 5.86e-11 | | *** |
| Deviant (vs Control) | 2.72 | 2.43 | 258.00 | 1.12 | 0.26390 | | |
| **Tukey post-hoc contrasts (Holm-corrected p-values)** | | | | | | | |
| Standard – Control | 16.60 | 2.43 | 6.84 | 6.84 | 2.44e-11 | 2.44e-11 | *** |
| Deviant – Control | 2.72 | 2.43 | 1.12 | 1.12 | 0.26290 | 0.26290 | |
| Deviant – Standard | -13.88 | 2.43 | -5.72 | -5.72 | 2.18e-08 | 2.18e-08 | *** |

##### Random Effects of the Linear Mixed-Effects Model

| Group | Name | Variance | Std. Deviation |
|---|---|---|---|
| Channel | Intercept | 1733.26 | 41.63 |
| Residual | NA | 383.43 | 19.58 |

Note: Model estimates and standard errors for fixed effects are shown first. Tukey's post-hoc pairwise comparisons (Holm-corrected p-values) are shown below the model estimates. Random effects report variance components for the random intercept. Significance stars: *p < 0.05; **p < 0.01; ***p < 0.001.

#### Supplementary Table 12

##### Early SSA and MMN Effects in Supragranular Single Units (Supplementary Figure 1H)

| Effect | Estimate | Std. Error / SEM | df / z | t/z value | p-value | Adj. p-value (Holm) | Sig. |
|---|---|---|---|---|---|---|---|
| Intercept | 3.31 | 1.11 | 78.00 | 2.98 | 0.00390 | | ** |
| Standard (vs Control) | -1.81 | 1.57 | 78.00 | -1.15 | 0.25310 | | |
| Deviant (vs Control) | 3.22 | 1.57 | 78.00 | 2.04 | 0.04450 | | * |
| **Tukey post-hoc contrasts (Holm-corrected p-values)** | | | | | | | |
| Standard – Control | -1.81 | 1.57 | -1.15 | -1.15 | 0.24960 | 0.24960 | |
| Deviant – Control | 3.22 | 1.57 | 2.04 | 2.04 | 0.08220 | 0.08220 | |
| Deviant – Standard | 5.03 | 1.57 | 3.19 | 3.19 | 0.00421 | 0.00421 | ** |

##### Random Effects of the Linear Mixed-Effects Model

| Group | Name | Variance | Std. Deviation |
|---|---|---|---|
| Neuron | Intercept | 0 | 0 |
| Residual | NA | 33.46 | 5.78 |

Note: Model estimates and standard errors for fixed effects are shown first. Tukey's post-hoc pairwise comparisons (Holm-corrected p-values) are shown below the model estimates. Random effects report variance components for the random intercept. Significance stars: *p < 0.05; **p < 0.01; ***p < 0.001.

#### Supplementary Table 13

##### Early SSA and MMN Effects in Granular Single Units (Supplementary Figure 1K)

| Effect | Estimate | Std. Error / SEM | df / z | t/z value | p-value | Adj. p-value (Holm) | Sig. |
|---|---|---|---|---|---|---|---|
| Intercept | 5.50 | 1.34 | 30.93 | 4.10 | 0.00028 | | *** |
| Standard (vs Control) | -3.40 | 1.25 | 34.00 | -2.72 | 0.01010 | | ** |
| Deviant (vs Control) | 1.47 | 1.25 | 34.00 | 1.18 | 0.24680 | | |
| **Tukey post-hoc contrasts (Holm-corrected p-values)** | | | | | | | |
| Standard – Control | -3.40 | 1.25 | -2.72 | -2.72 | 0.01287 | 0.01287 | ** |
| Deviant – Control | 1.47 | 1.25 | 1.18 | 1.18 | 0.23860 | 0.23860 | |
| Deviant – Standard | 4.87 | 1.25 | 3.90 | 3.90 | 0.00028 | 0.00028 | *** |

##### Random Effects of the Linear Mixed-Effects Model

| Group | Name | Variance | Std. Deviation |
|---|---|---|---|
| Neuron | Intercept | 18.51 | 4.30 |
| Residual | NA | 13.99 | 3.74 |

Note: Model estimates and standard errors for fixed effects are shown first. Tukey's post-hoc pairwise comparisons (Holm-corrected p-values) are shown below the model estimates. Random effects report variance components for the random intercept. Significance stars: *p < 0.05; **p < 0.01; ***p < 0.001.

#### Supplementary Table 14

##### Early SSA and MMN Effects in Infragranular Single Units (Supplementary Figure 1N)

| Effect | Estimate | Std. Error / SEM | df / z | t/z value | p-value | Adj. p-value (Holm) | Sig. |
|---|---|---|---|---|---|---|---|
| Intercept | 5.62 | 0.62 | 186.39 | 9.03 | 2.18e-16 | | *** |
| Standard (vs Control) | -4.01 | 0.73 | 150.00 | -5.53 | 1.39e-07 | | *** |
| Deviant (vs Control) | 1.69 | 0.73 | 150.00 | 2.33 | 0.02140 | | * |
| **Tukey post-hoc contrasts (Holm-corrected p-values)** | | | | | | | |
| Standard – Control | -4.01 | 0.73 | -5.53 | -5.53 | 6.42e-08 | 6.42e-08 | *** |
| Deviant – Control | 1.69 | 0.73 | 2.33 | 2.33 | 0.02000 | 0.02000 | * |
| Deviant – Standard | 5.70 | 0.73 | 7.86 | 7.86 | 1.20e-14 | 1.20e-14 | *** |

###### Random Effects of the Linear Mixed-Effects Model

| Group | Name | Variance | Std. Deviation |
|---|---|---|---|
| Neuron | Intercept | 9.48 | 3.08 |
| Residual | NA | 19.98 | 4.47 |

Note: Model estimates and standard errors for fixed effects are shown first. Tukey's post-hoc pairwise comparisons (Holm-corrected p-values) are shown below the model estimates. Random effects report variance components for the random intercept. Significance stars: *p < 0.05; **p < 0.01; ***p < 0.001.

#### Supplementary Table 15

##### Late DD and MMN Effects in Supragranular Single Units (Supplementary Figure 1I)

| Effect | Estimate | Std. Error / SEM | df / z | t/z value | p-value | Adj. p-value (Holm) | Sig. |
|---|---|---|---|---|---|---|---|
| Intercept | 4.45 | 1.51 | 46.83 | 2.94 | 0.00506 | | ** |
| Standard (vs Control) | 0.64 | 1.39 | 52.00 | 0.46 | 0.64550 | | |
| Deviant (vs Control) | 1.65 | 1.39 | 52.00 | 1.19 | 0.24080 | | |
| **Tukey post-hoc contrasts (Holm-corrected p-values)** | | | | | | | |
| Standard – Control | 0.64 | 1.39 | 0.46 | 0.46 | 0.93850 | 0.93850 | |
| Deviant – Control | 1.65 | 1.39 | 1.19 | 1.19 | 0.70630 | 0.70630 | |
| Deviant – Standard | 1.01 | 1.39 | 0.72 | 0.72 | 0.93850 | 0.93850 | |

##### Random Effects of the Linear Mixed-Effects Model

| Group | Name | Variance | Std. Deviation |
|---|---|---|---|
| Neuron | Intercept | 35.58 | 5.96 |
| Residual | NA | 26.10 | 5.11 |

Note: Model estimates and standard errors for fixed effects are shown first. Tukey's post-hoc pairwise comparisons (Holm-corrected p-values) are shown below the model estimates. Random effects report variance components for the random intercept. Significance stars: *p < 0.05; **p < 0.01; ***p < 0.001.

#### Supplementary Table 16

##### Late DD and MMN Effects in Granular Single Units (Supplementary Figure 1L)

| Effect | Estimate | Std. Error / SEM | df / z | t/z value | p-value | Adj. p-value (Holm) | Sig. |
|---|---|---|---|---|---|---|---|
| Intercept | 4.19 | 1.06 | 38.87 | 3.96 | 0.00031 | | *** |
| Standard (vs Control) | -0.90 | 1.17 | 34.00 | -0.77 | 0.44600 | | |
| Deviant (vs Control) | -0.19 | 1.17 | 34.00 | -0.16 | 0.87320 | | |
| **Tukey post-hoc contrasts (Holm-corrected p-values)** | | | | | | | |
| Standard – Control | -0.90 | 1.17 | -0.77 | -0.77 | 1 | 1 | |
| Deviant – Control | -0.19 | 1.17 | -0.16 | -0.16 | 1 | 1 | |
| Deviant – Standard | 0.71 | 1.17 | 0.61 | 0.61 | 1 | 1 | |

##### Random Effects of the Linear Mixed-Effects Model

| Group | Name | Variance | Std. Deviation |
|---|---|---|---|
| Neuron | Intercept | 7.99 | 2.83 |
| Residual | NA | 12.23 | 3.50 |

Note: Model estimates and standard errors for fixed effects are shown first. Tukey's post-hoc pairwise comparisons (Holm-corrected p-values) are shown below the model estimates. Random effects report variance components for the random intercept. Significance stars: *p < 0.05; **p < 0.01; ***p < 0.001.

#### Supplementary Table 17

##### Late DD and MMN Effects in Infragranular Single Units (Supplementary Figure 1O)

| Effect | Estimate | Std. Error / SEM | df / z | t/z value | p-value | Adj. p-value (Holm) | Sig. |
|---|---|---|---|---|---|---|---|
| Intercept | 5.42 | 0.59 | 163.78 | 9.17 | 1.95e-16 | | *** |
| Standard (vs Control) | -0.78 | 0.63 | 150.00 | -1.23 | 0.21910 | | |
| Deviant (vs Control) | 1.44 | 0.63 | 150.00 | 2.29 | 0.02320 | | * |
| **Tukey post-hoc contrasts (Holm-corrected p-values)** | | | | | | | |
| Standard – Control | -0.78 | 0.63 | -1.23 | -1.23 | 0.21720 | 0.21720 | |
| Deviant – Control | 1.44 | 0.63 | 2.29 | 2.29 | 0.04370 | 0.04370 | * |
| Deviant – Standard | 2.22 | 0.63 | 3.53 | 3.53 | 0.00126 | 0.00126 | ** |

##### Random Effects of the Linear Mixed-Effects Model

| Group | Name | Variance | Std. Deviation |
|---|---|---|---|
| Neuron | Intercept | 11.47 | 3.39 |
| Residual | NA | 15.07 | 3.88 |

Note: Model estimates and standard errors for fixed effects are shown first. Tukey's post-hoc pairwise comparisons (Holm-corrected p-values) are shown below the model estimates. Random effects report variance components for the random intercept. Significance stars: *p < 0.05; **p < 0.01; ***p < 0.001.

#### Supplementary Table 18

##### Decoding Accuracy Across Epochs and Conditions for Orientation Prediction (Figure 3A–C)

| Epoch/Condition | Mean Accuracy | 95% CI | p-value (perm test) | Effect Size (SMD) | Sig. |
|---|---|---|---|---|---|
| Baseline – Control | 0.603 | [0.424, 0.782] | 0.2525 | 1.14 | |
| Baseline – Standard | 0.720 | [0.573, 0.867] | 0.0029 | 2.97 | ** |
| Baseline – Deviant | 0.742 | [0.581, 0.903] | 0.0028 | 2.99 | ** |
| Early Change – Control | 0.141 | [-0.012, 0.294] | 0.0671 | 1.83 | |
| Early Change – Standard | 0.098 | [-0.037, 0.233] | 0.1495 | 1.44 | |
| Early Change – Deviant | 0.200 | [0.123, 0.277] | 0.0000 | 5.13 | *** |
| Late Change – Control | 0.305 | [0.206, 0.404] | 0.0000 | 6.10 | *** |
| Late Change – Standard | 0.149 | [0.044, 0.254] | 0.0049 | 2.81 | ** |
| Late Change – Deviant | 0.234 | [0.178, 0.290] | 0.0000 | 8.36 | *** |

Note: Mean decoding accuracy, 95% bootstrap confidence intervals, permutation p-values, and bias-corrected standardized mean differences (SMD) are reported for each condition and epoch. Significance: *$p < 0.05$; **$p < 0.01$; ***$p < 0.001$.

#### Supplementary Table 19

##### Levene's Test for Manhattan Distance Dissimilarity Across Contrasts (Figure 3E-F)

| Contrast | F-statistic | df1 | df2 | p-value | Sig. |
|---|---|---|---|---|---|
| Baseline: Standard–Deviant vs. Standard–Control | 25.287 | 1 | 278 | 0.0000 | *** |
| Baseline: Control–Standard vs. Control–Deviant | 0.069 | 1 | 278 | 0.7926 | |
| Baseline: Deviant–Standard vs. Deviant–Control | 23.774 | 1 | 278 | 0.0000 | *** |
| Standard: Baseline–Early vs. Deviant: Baseline–Early | 7.184 | 1 | 278 | 0.0078 | ** |

Note: Median-based Levene's test for equality of variances in population dissimilarity (Manhattan distance) across contrasts. Significance: *p < 0.05; **p < 0.01; ***p < 0.001.

#### Supplementary Table 20

##### Number of Orientation-Tuned Neurons per Trial Type and Epoch (Figure 4C)

| Category | Early Epoch | Late Epoch |
|---|---|---|
| **Significant in only one trial type** | | |
| Control only | 7 | 12 |
| Standard only | 20 | 23 |
| Deviant only | 11 | 24 |
| **Significant in two trial types** | | |
| Control & Standard | 4 | 6 |
| Standard & Deviant | 9 | 14 |
| Deviant & Control | 5 | 4 |
| **Significant in all three trial types** | | |
| All three trial types | 3 | 2 |
| **No significant OPI** | | |
| No significant OPI | 82 | 56 |

Note: Cell counts indicate neurons with significant orientation preference index (OPI, permutation $p < 0.05$) in the indicated trial type(s) and epoch. 'No significant OPI' indicates neurons with non-significant OPI in all conditions. For cells significant in two trial types, combinations are listed.

#### Supplementary Table 21

##### Chi-Square Tests for Preference Inversion Frequencies Across Trial Type Pairs (Figure 4D–H)

| Epoch/Condition Comparison | N Cells | N Switches | $\chi^2$ | p-value | Sig. |
|---|---|---|---|---|---|
| Early: Standard vs. Control | 7 | 1 | 1.08 | 0.2990 | |
| Early: Standard vs. Deviant | 12 | 5 | 6.32 | 0.0120 | * |
| Early: Control vs. Deviant | 8 | 1 | 1.07 | 0.3020 | |
| Late: Standard vs. Control | 8 | 2 | 2.29 | 0.1310 | |
| Late: Standard vs. Deviant | 16 | 16 | 7.38 | 0.0070 | ** |
| Late: Control vs. Deviant | 6 | 2 | 2.40 | 0.1210 | |

Note: Proportions of neurons exhibiting orientation preference inversion (OPI switch) across trial type pairs and epochs. p-values from chi-square tests. Significance: *$p < 0.05$; **$p < 0.01$; ***$p < 0.001$.

#### Supplementary Table 22

##### Decoder Accuracy Comparisons Between Preference-Switching and Stable Neuron Subsets Across Trial Types and Epochs (Early, Figure 5A–C)

| Trial Type | Mean Accuracy | 95% CI | p-value | Effect Size (SMD) | Sig. |
|---|---|---|---|---|---|
| Control | -0.001 | [-1.493, 1.492] | 0.9995 | -0.00 | |
| Standard | 0.010 | [-1.586, 1.607] | 0.9897 | 0.01 | |
| Deviant | 0.042 | [-1.797, 1.881] | 0.9641 | 0.04 | |

Note: Mean decoding accuracy, 95% bootstrap confidence intervals, permutation p-values, and standardized mean differences (SMD) are reported for each trial type. Comparison is between preference-switching and matched stable neuron subsets. Significance: *p < 0.05; **p < 0.01; ***p < 0.001.

#### Supplementary Table 23

##### Decoder Accuracy Comparisons Between Preference-Switching and Stable Neuron Subsets Across Trial Types and Epochs (Late, Figure 5D–F)

| Trial Type | Mean Accuracy | 95% CI | p-value | Effect Size (SMD) | Sig. |
|---|---|---|---|---|---|
| Control | 0.001 | [-1.792, 1.794] | 0.9989 | 0.00 | |
| Standard | 0.043 | [-1.621, 1.706] | 0.9596 | 0.05 | |
| Deviant | 0.015 | [-1.883, 1.913] | 0.9874 | 0.02 | |

Note: Mean decoding accuracy, 95% bootstrap confidence intervals, permutation p-values, and standardized mean differences (SMD) are reported for each trial type. Comparison is between preference-switching and matched stable neuron subsets. Significance: *p < 0.05; **p < 0.01; ***p < 0.001.

#### Supplementary Table 24

##### Decoder Accuracy for Preference-Switching and Stable Neuron Subsets (Switchers-Only Analysis, Early, Supplementary Figure 5A–C)

| Trial Type | Mean Accuracy | 95% CI | p-value | Effect Size (SMD) | Sig. |
|---|---|---|---|---|---|
| Control | -0.178 | [-1.477, 1.120] | 0.7864 | -0.27 | |
| Standard | 0.029 | [-1.211, 1.269] | 0.9629 | 0.05 | |
| Deviant | 0.110 | [-1.534, 1.754] | 0.8953 | 0.13 | |

Note: Mean decoding accuracy, 95% bootstrap confidence intervals, permutation p-values, and standardized mean differences (SMD) are reported for each trial type. Analysis includes only preference-switching and matched stable neuron subsets (switchers-only analysis). Significance: *p < 0.05; **p < 0.01; ***p < 0.001.

#### Supplementary Table 25

##### Decoder Accuracy for Preference-Switching and Stable Neuron Subsets (Switchers-Only Analysis, Late, Supplementary Figure 5D–F)

| Trial Type | Mean Accuracy | 95% CI | p-value | Effect Size (SMD) | Sig. |
|---|---|---|---|---|---|
| Control | 0.189 | [-1.141, 1.519] | 0.7796 | 0.28 | |
| Standard | 0.311 | [-1.040, 1.661] | 0.6500 | 0.45 | |
| Deviant | 0.020 | [-1.746, 1.786] | 0.9824 | 0.02 | |

Note: Mean decoding accuracy, 95% bootstrap confidence intervals, permutation p-values, and standardized mean differences (SMD) are reported for each trial type. Analysis includes only preference-switching and matched stable neuron subsets (switchers-only analysis). Significance: *p < 0.05; **p < 0.01; ***p < 0.001.

Only the trial type and epoch combinations relevant to each figure panel are shown for Tables 22–25. Additional comparisons (all condition and epoch combinations) are available in the supplementary source data file or upon request.

#### Supplementary Table 26

##### Summary of Included Animals and Sessions Used for Analysis (Methods Section)

| Animal | Session | N Electrodes | N Units (Total) | N Units (Responsive) |
|---|---|---|---|---|
| 1.52 | 1.52Rec1 | 18 | 13 | 13 |
| 1.57 | 1.57Rec1 | 20 | 29 | 21 |
| 1.59 | 1.59Rec1 | 20 | 7 | 6 |
| | 1.59Rec3 | 20 | 0 | 0 |
| 1.6 | 1.60Rec1 | 18 | 0 | 0 |
| 1.61 | 1.61Rec4 | 20 | 2 | 2 |
| 1.62 | 1.62Rec4 | 18 | 26 | 25 |
| 1.63 | 1.63Rec1 | 20 | 4 | 3 |
| | 1.63Rec3 | 20 | 17 | 17 |
| | 1.63Rec4 | 20 | 20 | 15 |
| 1.64 | 1.64Rec1 | 20 | 1 | 1 |
| | 1.64Rec3 | 18 | 10 | 9 |
| | 1.64Rec4 | 20 | 12 | 9 |

Note: Each row details an individual animal and recording session included in the analysis. N Units (Total) refers to the total number of sorted units per session; N Units (Responsive) refers to units that showed a significant response to sensory stimuli.
